# Spurious intragenic transcription is a hallmark of mammalian cellular senescence and tissue aging

**DOI:** 10.1101/2022.05.20.492816

**Authors:** P. Sen, G. Donahue, C. Li, Y. Lan, G. Egervari, N. Robertson, P. P. Shah, E. Kerkhoven, D. C. Schultz, P. D. Adams, S. L. Berger

## Abstract

Mammalian aging is characterized by the progressive loss of tissue integrity and function manifesting in ill health and increased risk for developing multiple chronic conditions. Accumulation of senescent cells in aging tissues partly contributes to this decline and targeted depletion of senescent cells *in vivo* ameliorates many age-related phenotypes. However, the fundamental molecular mechanisms responsible for the decline of cellular health and fitness during senescence and aging are largely unknown. In this study, we investigated whether chromatin-mediated loss of transcriptional fidelity, known to contribute to fitness and survival in yeast and worms, also occurs during human cellular senescence and mouse aging. Our findings reveal that aberrant transcription initiation inside genes is widespread in senescence and aging. It co-occurs with changes in the chromatin landscape and formation of non-canonical transcription start sites. Interventions that alter spurious transcripts have dramatic consequences on cellular health primarily affecting intracellular signal transduction pathways. We propose that spurious transcription is a conserved hallmark of aging that promotes a noisy transcriptome and degradation of coherent transcriptional networks.

---

In eukaryotes, the transcription of the genome into precursor mRNA is catalyzed by RNA polymerase II^1^. During RNA polymerase II transcription, repressive closed chromatin over the gene body and promoter is opened to allow for processive RNA polymerase-mediated transcription elongation. However, after the passage of RNA polymerase II it is crucial that the initial chromatin structure is restored to preserve genome integrity. A failure to do so leads to perturbances in nucleosome organization, exposure of intragenic (or cryptic) promoter-like sequences, and inappropriate initiation of aberrant transcripts from inside gene bodies^2^. Our previous work investigating aging in yeast and worms demonstrated a role for the gene body protective trimethylation mark on lysine 36 of histone H3 (H3K36me3) in preventing age-related intragenic cryptic transcription^3^. More recent studies of normal growth in mouse embryonic stem cells shows that gene-body DNA methylation co-operates with H3K36 methylation to prevent intragenic cryptic transcription^4^. Furthermore, chemotherapeutic agents such as DNA methyltransferase inhibitors and histone deacetylase inhibitors upregulate new cryptic transcription from unannotated intronic transcription starts sites (TSSs), which result in 5′-truncated open reading frames^5^. These findings suggest that tightly regulated epigenetic crosstalk exerted by histone methylation, histone acetylation and DNA methylation ensure reliable and faithful initiation of gene transcription.

Senescence is a state of stable cell proliferation arrest caused by a variety of cellular stressors, including DNA damage, activation of oncogenes, replicative exhaustion and other toxins^6,7^. Senescence functions to prevent cancer by halting damaged or precancerous cells from dividing^8,9^. Additionally, senescent cells secrete a unique profile of cytokines and other soluble factors collectively called the Senescence Associated Secretory Phenotype (SASP) that is thought to promote immune clearance of premalignant senescent cells^10^. However, chronic exposure to these same inflammatory cytokines can cause tissue damage and ultimately aging. Indeed, targeted elimination of senescent cells improves lifespan and healthspan in mice^11-13^. Thus, while senescence protects against cancer, it is also an important contributor to organismal aging.

Senescent cells show profound alterations in nuclear architecture and chromatin organization that are predicted to culminate in gene expression changes^14,15^. In support, investigation of transcriptome profiles in senescent cells have shown expression changes such as upregulation of inhibitors of cell cycle^16^ and SASP^10^, and downregulation of lamin B1^17-20^ and canonical histone proteins^21^. However, the exact relationship between chromatin dysregulation during senescence and aging and an altered transcriptome is less understood.

Increased variability and “noisy” transcription are observed features of aging^22^. Furthermore, there is a major divergence of the transcriptome and proteome with aging suggesting decoupling mechanisms at play^23^. Our results in this study identify a novel mechanism that contributes to noise in an aging transcriptome. The noise arises from a largely open chromatin structure over genes and loss of faithful transcription initiation in senescent cells and aging tissues. Our work indicates that this aberrant mechanism is conserved from yeast to humans and uncover a progressive loss of information content in transcriptional networks during aging.

## Results

### Genome-wide precision run-on mapping reveals many intragenic cryptic initiation sites

Given our previous observations in aging invertebrate systems^3^ and the recent discovery of intragenic cryptic transcription in mammalian cells^4,5,24^, we undertook an investigation of transcriptional fidelity and cryptic transcription in mammalian cellular senescence and aging. However, whereas yeast and worms have simple transcription patterns, mammalian gene units are complex with alternate start sites at the 5’ end, elaborate intron-exon organization, frequent alternative splicing and alternative poly-adenylation sites, as well as overlapping genes. Consequently, traditional RNA-seq approaches will not distinguish new TSS formation from other events. Given this obstacle, we instead set out to directly identify newly formed active TSSs, by employing precision run-on RNA sequencing methodologies (PRO-seq and PRO-cap ^25^). To our knowledge, this is the first genome-wide nascent transcription mapping in replicative senescent cells. We used lung fibroblast IMR90 cells that had reached senescence as a result of replicative exhaustion^26^. Proliferating cells were harvested at population doubling (PD) 20-30 and senescent cells were harvested at PD 70-80 after they had growth arrested and had upregulated markers of senescence such as beta-galactosidase (data not shown, Fig. S1.1A). PRO-seq and PRO-cap were performed with two independent cell populations and showed a strong correlation in peak signals detected by HOMER with FDR <0.001 (Fig. S1.1B-D). PRO-cap tags identify the precise initiation sites of capped (and therefore stable and mature) nascent transcripts, hence we used this peak list to perform all downstream analyses. We detected in both proliferating and senescent cells ∼116-122K PRO-cap peaks distributed in promoters, genic and intergenic regions (Fig. 1A, first two pie charts). We then imposed two stringent criteria for the identification of unique PRO-cap peaks in proliferating and senescent cells; a reads per million (rpm) cutoff of >4 to eliminate noise, and a fold change (FC) cutoff of >10 to identify peaks present in only one condition (Fig. S1.1E, #1). We note that the >4 rpm cutoff was qualitatively chosen after visual examination of peaks on the UCSC Genome Browser. Additionally, we required that there be no overlap between peaks in proliferating and senescent cells. A compartment analysis of these unique peaks in both proliferating (Unique Pro) and senescent (Unique Sen) cells showed a redistribution of peaks to genic (∼12K and ∼9.3K peaks in proliferating and senescent cells respectively) and intergenic (∼11K and ∼6.2K peaks in proliferating and senescent cells respectively) regions with major losses at promoters (Fig. 1A, last two pie charts and Fig. 1B). The percentage of unique genic peaks was greater in senescent cells compared to proliferating cells (58% vs 46% respectively), indicating that gene-internal cryptic initiation sites are proportionally more frequent in senescence.

**Fig. 1:**
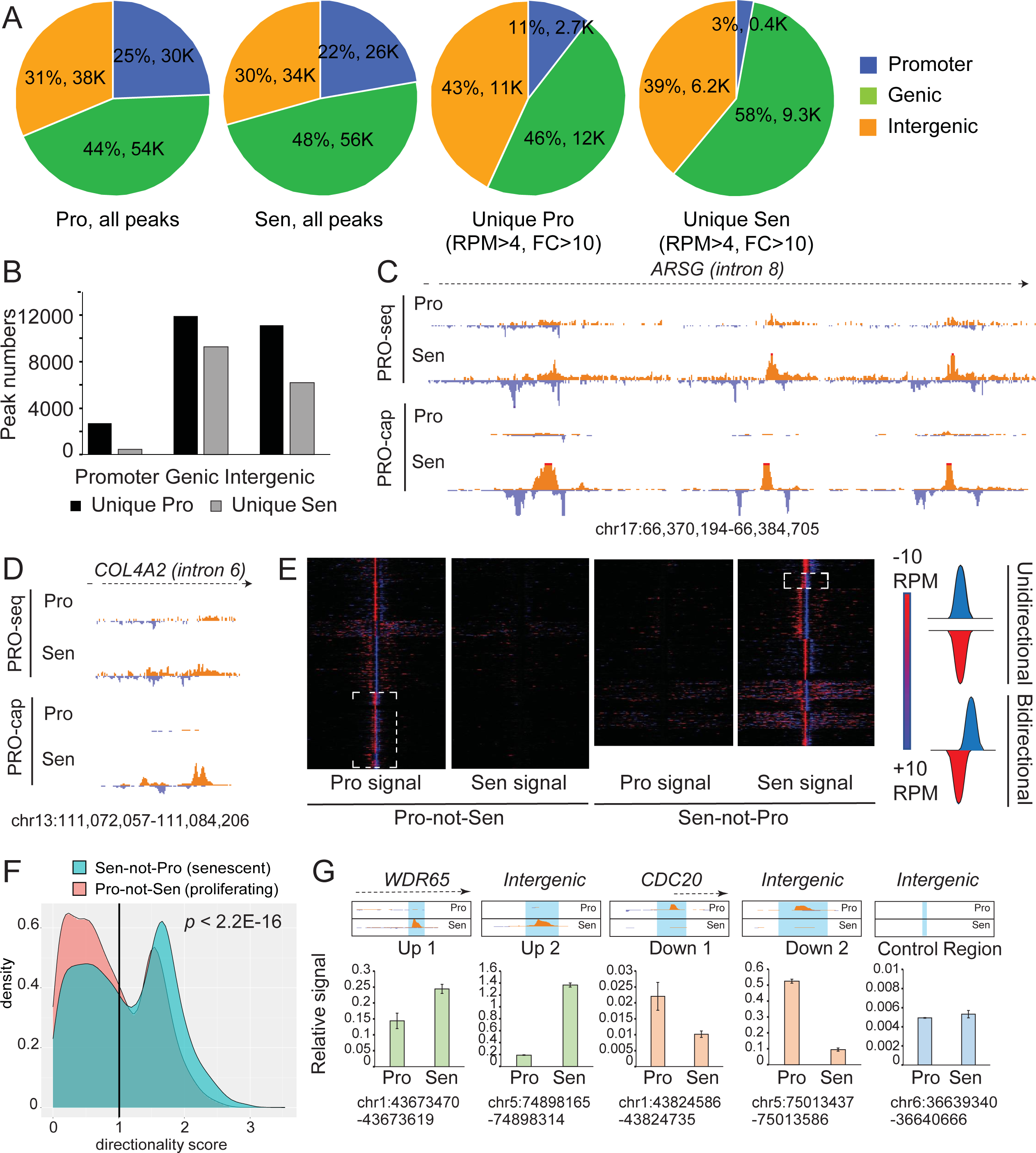
Cryptic TSSs are activated in proliferating and senescent cells. **(A)** Compartment analysis of all (first two pie charts) and unique (last two pie charts) PRO-cap peaks in proliferating and senescent cells as called by HOMER. Unique peaks were filtered to retain those that had a signal above 4rpm and a 10-fold enrichment by an AUC measurement. Promoters were assigned as 1 Kb around the TSS, genic as everything in the gene except the promoter region and intergenic as everything but promoter and gene. Proportions of peaks in a specific compartment are denoted as % and total numbers of peaks are indicated. **(B)** Peak numbers from the last two pie charts are plotted as a bar graph. **(C-D)** Browser track views of PRO-cap peaks detected in senescence at two gene-internal locations. Chromosomal coordinates are indicated below the browser view. **(E)** Heatmaps of genes that show evidence of cryptic transcription in proliferating and senescence conditions, centered around the unique PRO-cap peak. Blue and red indicate signal in plus and minus strands respectively. White dashed boxes show PRO-cap peaks with bidirectional transcription initiation. **(F)** Histogram showing the distribution of directionality scores of cryptic TSSs in proliferating and senescent cells. The vertical line represents a directionality score of 1 below which TSSs are classified as bidirectional. *p-*value is reported based on a coin library permutation test. **(G)** qPCR measurement of PRO-cap signal in senescent cells (Up), proliferating cells (Down) or a negative control region (Control Region). Error bars are from standard deviations of qPCR technical replicates.

To unambiguously identify cryptic TSSs inside gene bodies, we further filtered the unique PRO-cap peaks on two criteria. First, to distinguish new cryptic start sites from new canonical promoter-driven transcription, we chose peaks localizing to the 3’ end of the gene and at least 3.5 Kb away from any other annotated TSSs. Second, the peak list was filtered to remove GENCODE TSSs that might represent non-coding RNA start sites (Fig. S1.1E, #2 and #3). We selected a distance cutoff of 3.5 Kb because our analysis shows that two thirds of all ENCODE-annotated strong and weak promoters are within 3.5 Kb from their nearest TSS (Fig. S1.1F)^27^, thus, ensuring that TSSs from overlapping genes were removed. These criteria identified ∼4.8 and ∼4.2K novel intragenic cryptic sites in proliferating (“Pro-not-Sen”) and senescent (“Sen-not- Pro”) cells respectively (Table S1). Figs. 1C and 1D show two examples of typical cryptic TSSs detected in senescent cells within introns of genes *ARSG* (Arylsulfatase G) and *COL4A2* (collagen type IV alpha 2 chain). Extensive comparison of PRO-seq and PRO-cap tracks confirmed that our bioinformatic pipeline accurately detected new cryptic TSSs. A strand-specific heatmap surrounding cryptic TSSs shows a strong enrichment of PRO-cap tags in either the proliferating or senescent state compared to lack of signal at the same sites in the other condition (Fig. 1E). We noted that in the proliferating condition, ∼30% of the cryptic TSSs showed bidirectional transcription typical of enhancers, compared to only 8% in senescent cells (Fig. 1E dotted boxes). The proportion of bidirectional transcription was further confirmed by assigning a directionality score (see Methods) to transcripts from Pro-not-Sen and Sen-not-Pro cryptic TSSs based on a previous publication^28^. A histogram plot of cryptic transcript directionality score distribution in a 300bp region showed that a significant proportion of Pro-not-Sen cryptic transcripts had low directionality score in the proliferating condition, in other words, were bidirectional (Fig. 1F). Finally, we validated our sequencing data using the conventional qPCR approach by amplifying a region surrounding a PRO-cap peak enriched in senescent cells (Up), enriched in proliferating cells (Down) or a negative control region (Control Region) (Fig. 1G). Taken together, this initial survey of PRO-cap peaks revealed many cryptic TSSs in both proliferating and senescent cells, with a larger number of enhancer-like peaks in proliferating cells.

### Cryptic TSSs are more abundant in long genes with long introns

We next investigated the general features of genes that were prone to intragenic cryptic transcription. Genes with internal initiation sites were significantly longer than the average gene in the human genome (Fig. S1.2A), a feature that was previously identified in yeast and worms^29^. Additionally, these genes have fewer exons (Fig. S1.2B), fewer introns (Fig. S1.2C), shorter exons (Fig. S1.2D), longer introns (S1.2E), as well as shorter 5’ (Fig. S1.2F) and 3’ UTRs (Fig. S1.2G). These features were indistinguishable in the genes harboring cryptic TSSs in either proliferating or senescent conditions, suggesting that these general genic features show a high propensity for cryptic initiation regardless of cell state. We speculate that the vast excess of intronic sequence in these genes leads to transcription delays, increased RNA pol II residence time, and, hence, cryptic initiation^30^.

### Cryptic TSSs in proliferating cells are enhancer-like whereas those in senescent cells are promoter-like

Canonical TSSs have distinct histone post-translational modifications (hPTMs) and histone variants present in their vicinity^31,32^. We assayed the chromatin landscape around the cryptic TSSs to compare to canonical counterparts. We profiled chromatin accessibility by ATAC-seq^33^, multiple hPTMs, both acetylation and methylation sites in histone H3 and H4, and histone variants (H3.3 and H2A.Z) by ChIP-seq^34^. Genome-wide datasets generated in this study are detailed in Table S2. Averaged metaplots centered around cryptic PRO-cap peaks were constructed to display differential enrichments of the various histone modifications and chromatin accessibility. The marks in proliferating cells are traced with a thin line while those in senescent cells with a thick line. The central nucleosome showed modest accessibility by ATAC-seq in the proliferating state in both Pro-not-Sen and Sen-not-Pro peaks (Fig. 2A). However, in senescent cells, regardless of transcriptional activity, chromatin accessibility increased dramatically suggesting that cryptic TSSs are more open in the senescent state (Fig. 2A thick lines). In line with chromatin opening, there was higher enrichment of most histone acetylation marks such as H3K18ac (Fig. 2B), H3K23ac (Fig. 2C), H3K122ac (Fig. 2E) and H4K5ac (Fig. 2F) in senescence. H3K27ac was the only acetylation mark that increased with the direction of transcription, going up in the Pro-not-Sen sites in proliferation, and in the Sen-not-Pro sites in senescence (Fig. 2D). Correspondingly, the direction of occupancy of cognate histone acetyltransferases p300 (Fig. 2G) and CBP (Fig. 2H) reflected the enrichment of H3K27ac.

**Fig. 2:**
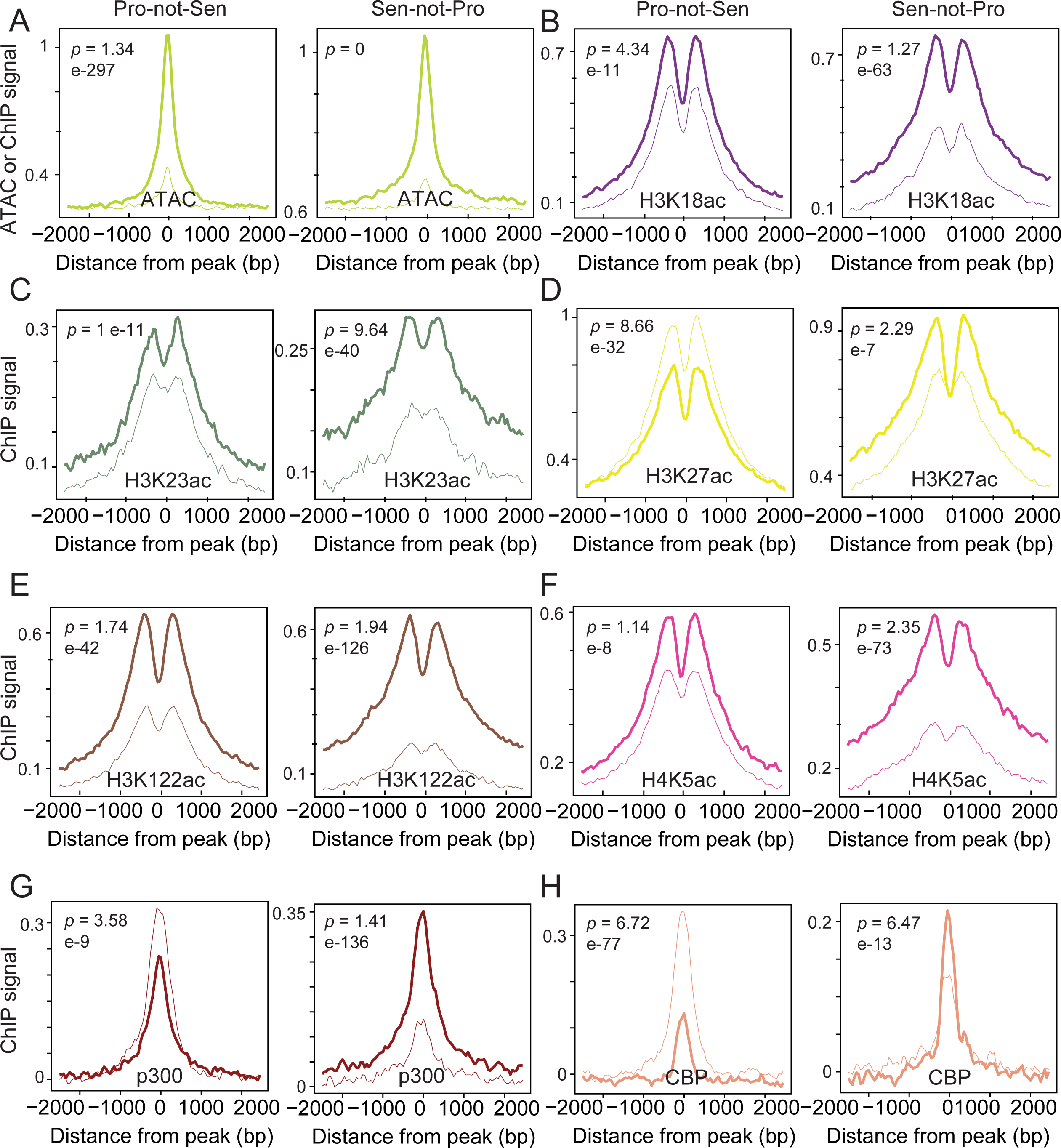
Chromatin acetylation profiles around cryptic TSSs suggest H3K27ac drive cryptic transcription Metaplots of ATAC-seq. **(A)**, H3K18ac **(B)**, H3K23ac **(C)**, H3K27ac **(D)**, H3K122ac **(E)**, H4K5ac **(F)**, p300 **(G)**, and CBP **(H)** signal are shown in a 2 Kb region surrounding cryptic TSSs. Thin lines represent proliferating cells and thick lines senescent cells. *p-*values are reported based on a Mann-Whitney U test.

In contrast to histone acetylation, cryptic TSSs were devoid of certain histone methylation such as H3K27me3 (Fig.S3.2A), H3K36me3 (Fig. S3.2B), H3K79me1 (Fig. S3.2C) and H3K79me2 (Fig. S3.2D).

However, H3K4me1 and H3K4me3 levels at the cryptic TSSs showed interesting changes. The ratio of these two modifications determines whether a regulatory region is classified as an enhancer (high H3K4me1/H3K4me3) or promoter (high H3K4me3/H3K4me1)^35^. The Pro-not-Sen sites showed considerable amounts of both modifications in the proliferating state with a loss of enrichment in the senescent state (Fig. 3A-B), suggesting that Pro-not-Sen sites are likely active promoters and enhancers that are delicensed in senescence. Given the high ratio of H3K4me1/H3K4me3 at these Pro-not-Sen sites and the fact that we filtered out putative TSSs from overlapping genes, we conclude that the majority of Pro-not-Sen sites represent active enhancers that become delicensed in senescence. The Sen-not-Pro sites showed a distinct pattern, gaining significant H3K4me3 at these sites while losing H3K4me1 (Fig. 3A-B), suggesting these sites are poised enhancers in proliferating cells that undergo activation to a promoter-like state in senescence. Our results indicate that enhancer-to-promoter conversion is a frequent event in senescent cells.

**Fig. 3:**
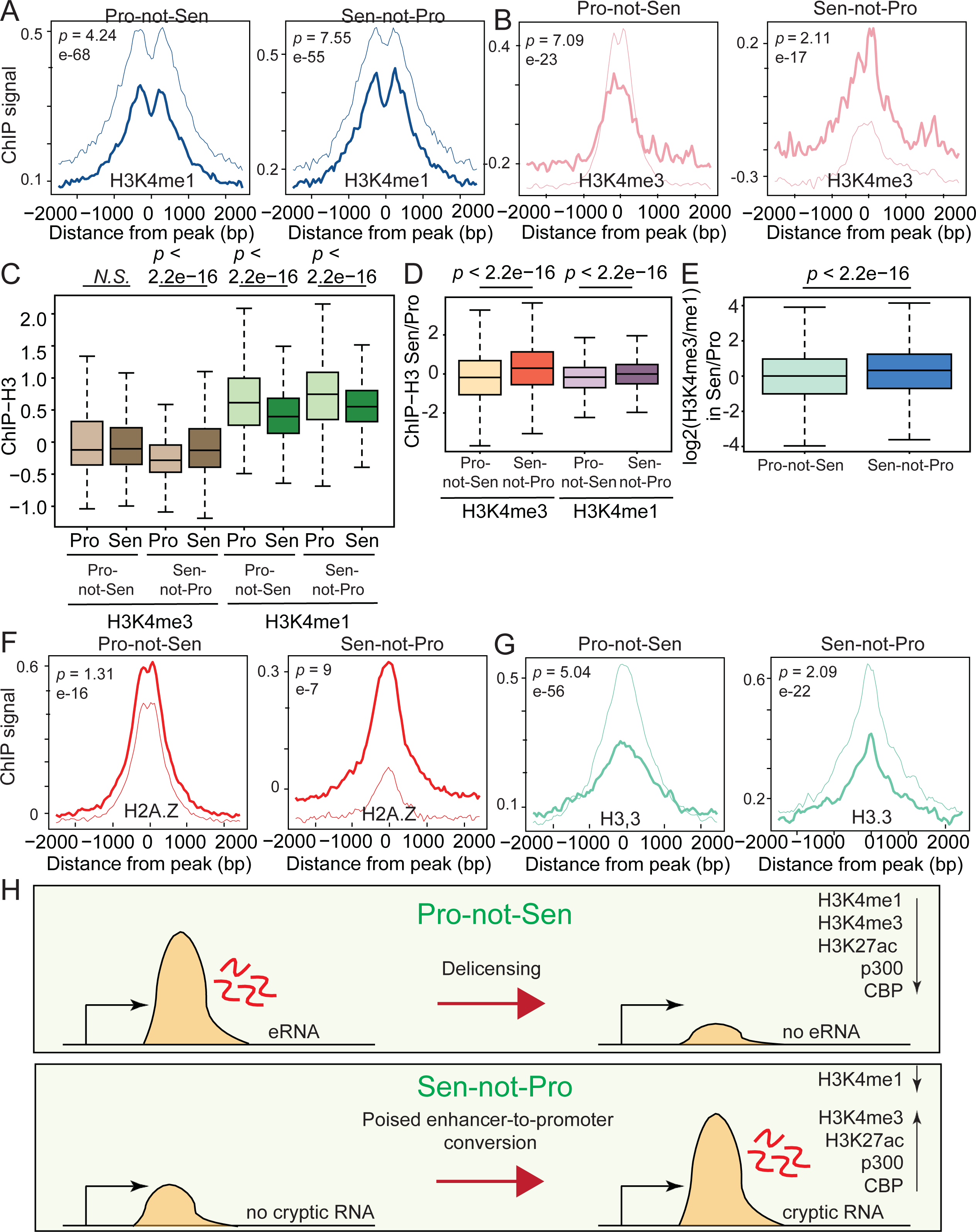
Chromatin methylation profiles around cryptic TSSs suggest an enhancer-to-promoter conversion in senescent cells. **(A-B)** Metaplots of H3K4me1 **(A)**, and H3K4me3 **(B)** signal are shown in a 2 Kb region surrounding cryptic TSSs. **(C)** Boxplot showing the H3 normalized H3K4me1 and H3K4me3 ChIP signal individually at Pro-not-Sen and Sen-not-Pro cryptic sites in proliferating and senescent cells. **(D)** Boxplot showing the gain in H3K4me3 in senescent cells at Sen-not-Pro cryptic sites. **(E)** Boxplot showing the enhancer-to-promoter conversion plotted as a ratio of ratios of H3K4me3/H3K4me1 signal in senescent vs proliferating cells. **(F-G)** Metaplots of H2A.Z **(F)** and H3.3 **(G)** are shown in a 2 Kb region surrounding cryptic TSSs. In the metaplots (A-B and F-G), the thin lines represent proliferating cells and thick lines senescent cells. *p-*values are reported based on a Mann-Whitney U test. **(H)** Illustration of the chromatin changes occurring at Pro-not-Sen and Sen-not-Pro sites.

To better depict the dynamic changes in H3K4me1 and H3K4me3 around cryptic sites, we box plotted these modifications individually (Fig. 3C) as well as in ratios (Fig. 3D-E) to show the extent of delicensing or promoter conversion. Fig. 3C shows that in the Sen-not-Pro cryptic sites there is a substantial increase of H3K4me3 in senescent cells while H3K4me1 decreased. The state of these modifications in senescent and proliferating cells is depicted as a ratio (Senescent vs Proliferating) in Fig. 3D which shows a positive signal for H3K4me3 (orange box) at Sen-not-Pro sites. Finally, in Fig. 3E, we represent a ratio of ratios, plotting the H3K4me3/H3K4me1 ratio in Senescent versus Proliferating condition, again showing a positive signal at Sen- not-Pro sites.

We next noted that cryptic TSSs became marked with a well-positioned H2A.Z while losing H3.3 occupancy in senescence (Fig. 3F-G). The enrichment of H2A.Z at Sen-not-Pro sites was particularly prominent and reminiscent of well-known H2A.Z occupancy near canonical TSSs^36^. The metaplot profiles show that the nucleosomes flanking the central PRO-cap peak (-1 and +1) were enriched in H3K4me3 (Fig. 3B) while the -2 and +2 nucleosomes were decorated with H3K4me1 (Fig. 3A) and histone acetylation (Fig. 2B-F) . These data suggest dynamic chromatin restructuring at cryptic TSSs. Overall, the chromatin profiling indicated that Pro-not-Sen sites are intragenic enhancers that become delicensed in senescence, while Sen-not-Pro sites are sites of enhancer-to-promoter conversions (Fig. 3H).

### Canonical and cryptic TSSs share some common features but also show distinct chromatin patterns

To directly compare cryptic TSSs to canonical TSSs, we profiled the same chromatin marks at the annotated TSSs of the cryptic genes. This comparison revealed important similarities and conspicuous differences in chromatin composition (Figs. 2, 3, S2.1, S3.1 and S3.2). For example, both canonical and cryptic TSSs were depleted of H3K36me3 (Figs. S3.1B and S3.2B) despite that cryptic TSSs are located within the gene body where H3K36me3 levels are usually high^37^. Canonical TSSs contain H3.3 and H2A.Z within one nucleosome (as reported^38^) occurring on both sides of the TSS; cryptic TSSs in contrast are directly occupied by an H2A.Z/H3.3-containing nucleosome suggesting that normally cryptic sites are protected. The enrichment of these variants despite the presence of transcripts could reflect cell-to-cell heterogeneity whereby some cells that initiate cryptic transcription do not have nucleosomes at that site whereas other cells have not yet evicted the nucleosome. Furthermore, in canonical TSSs, most histone modifications and variants are positioned at -1 and +1 nucleosomes (Fig. S2.1B-F and S3.1F-H). While H3K4me1 in cryptic TSSs is restricted to the -2 and +2 nucleosomes (Fig. 3A), H3K4me1 is substantially depleted in the region spanning -2 to +2 on canonical promoters (Fig. S3.1E). These differences suggest that nucleosomes surrounding a cryptic TSS exhibit “fuzzier” positioning compared to canonical TSSs.

### Genes harboring cryptic TSSs are actively expressed

To test whether cryptic transcription interfered with full length transcription from the cognate canonical TSS, we assessed the correlation of steady state RNA-seq or nascent PRO-seq signals at canonical TSSs to PRO-cap signal at the cryptic sites in the same genes. Signal differences were measured by AUC (Area Under the Curve) analysis which quantifies a normalized tag density over a region in the genome (here the PRO-cap peak at the cryptic TSS). PRO-cap signals at cryptic sites were divided into deciles and the PRO-seq signal was measured in each decile; we found concordant transcription strength, showing a strong correlation between PRO-cap and PRO-seq (Fig. S3.3A-B, statistical tests in Table S4). While there was no correlation between PRO-cap signal amplitude and steady state RNA levels, PRO-cap cryptic peaks were formed in actively expressed genes in either condition (Fig. S3.3C-D, note the positive y-axis across the deciles, statistical tests in Table S4). A similar pattern was observed with nascent transcription with occurrence of PRO-cap peaks at genes that have high PRO-seq signal at the TSS (Fig. S3.3E-F, statistical tests in Table S4). These experiments indicate that cryptic transcription is coincident with ordinary gene transcription and nascent transcription, occurring primarily in active genes. Cryptic transcription does not appear to interfere with canonical transcription in the cognate gene.

### Cryptic TSSs show loss of DNA and histone methylation in senescence

Recent reports that investigate spurious intragenic transcription in mammalian systems suggest the major cause of its upregulation is loss of gene-body DNA and H3K36 methylation and increase of histone acetylation. These studies employed shRNA-mediated depletion or drug inhibition of DNA methyltransferase 3B or SET domain containing 2 proteins (DNMT3B/SETD2) or administration of histone deacetylase (HDAC) inhibitors to reduce methylation and increase acetylation respectively^4,5^. Given the close relationship between DNA/histone methylation and cryptic transcription, we investigated the DNA methylation status directly under a new cryptic site as well as the entire gene. We noted that the proportion of CG methylation was low (∼30%, Fig. 4A 500bp around) in Pro-not-Sen sites supporting the notion that cryptic transcripts originate at regions of low methylation and that enhancers are generally hypomethylated. A parameter sweep (125bp, 250bp, 500bp or 1000bp) downstream of the cryptic site showed a progressive increase in the methylation signal. Importantly, the methylation state did not change at these Pro-not-Sen sites in senescent cells (Fig. 4A). In contrast, a similar analysis of DNA methylation status at Sen-not-Pro sites revealed a significant loss of DNA methylation in senescence suggesting an active demethylation event at these specific sites that promotes internal initiation (Fig. 4B). To study the overall methylation status of cryptic genes we generated metaplots of DNA methylation and H3K36me3 over the gene bodies. To our surprise we found that DNA CG methylation was high in both Pro-not-Sen and Sen-not-Pro genes compared to a random set of background genes, reflective of their high transcriptional activity (Fig. 4C-F). Additionally, we noted significantly higher levels of H3K36me3 levels at Sen- not-Pro genes in senescence, again suggestive of higher RNA pol II activity at these genes. Taken together, the transcriptional levels and DNA/histone methylation distribution at cryptic genes suggest that cryptic transcription may be a consequence of polymerase “overloading” at these genes with then increased propensity to initiate transcription from hypomethylated aberrant locations in the context of an open and unstable senescent epigenome. While this data contradicts previous observations of an inverse relationship between DNA/histone methylation on cryptic transcription^4,5^, our results show that *local* rather than *total* gene body changes in chromatin marks drive the formation of cryptic TSSs.

**Fig. 4:**
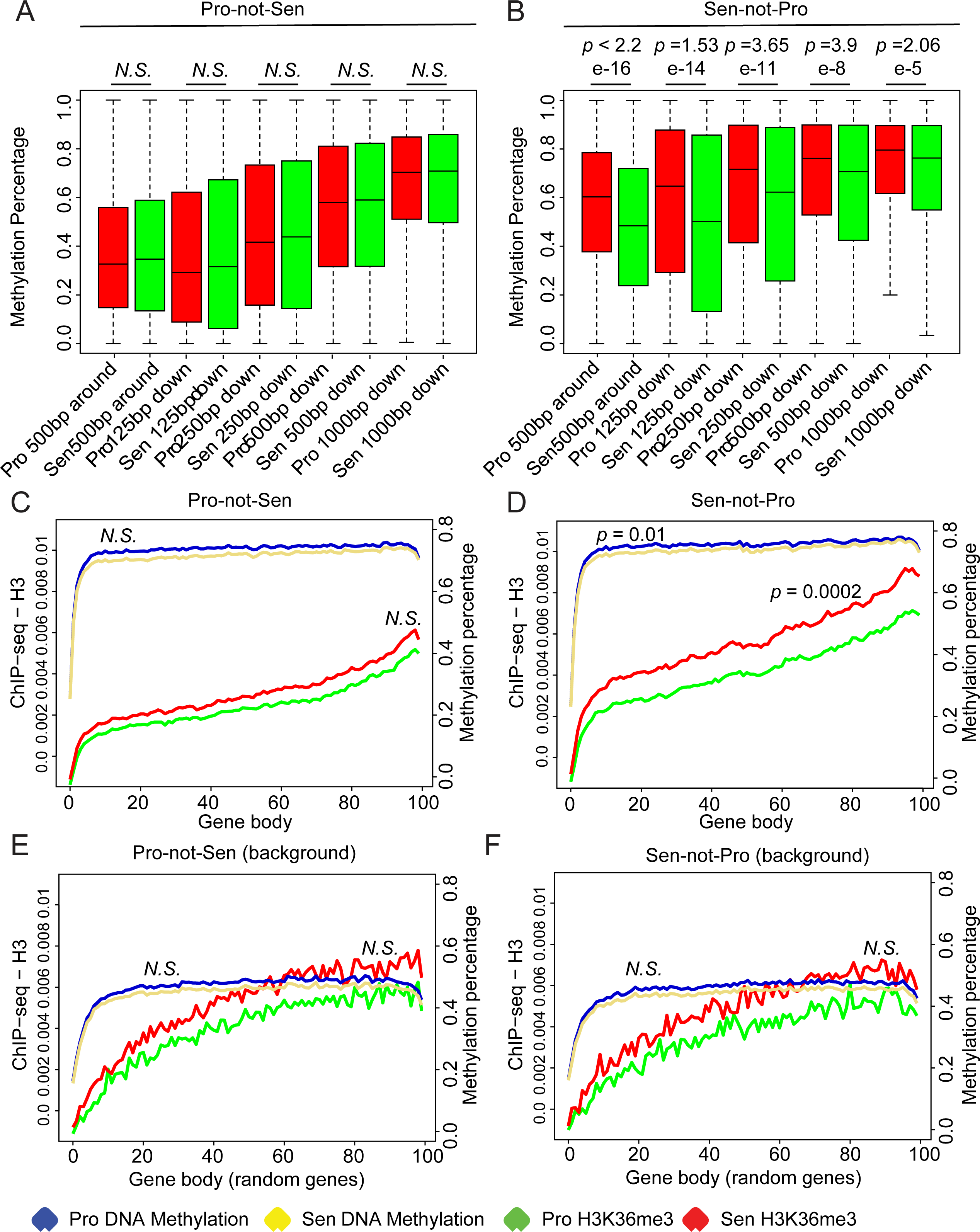
Sen-not-Pro cryptic TSSs lose DNA methylation in senescence. **(A-B)** Parameter sweep of DNA methylation percentage at and around Pro-not-Sen **(A)** or Sen-not-Pro **(B)** cryptic TSSs in proliferating and senescent cells. **(C-D)** Metaplots of DNA methylation, and normalized H3K36me3 signal over Pro-not-Sen **(C)** or Sen-not-Pro **(D)** cryptic genes or an equal number of random genes **(E-F)**. *p-*values are reported based on a Mann-Whitney U test. *N.S.* indicates not significant.

### AP-1 family of transcription factors drive cryptic transcription

For insight into biological pathways affected by cryptic transcription, we performed Gene Ontology (GO) analysis of the genes containing unique cryptic sites in proliferating and senescent cells. GO terms associated with the cryptic genes were linked to membrane and cytoplasmic components, GTPase activity, cytoskeleton, cell shape/adhesion and signal transduction pathways (Fig. S4.1A-B). There were no marked distinctions in Gene Ontology between the two classes of cryptic genes suggesting that in general, cryptic gene features (such as gene length, intron length, high transcriptional activity etc.) or specific recruitment of transcription factors (TFs) rather than functional ontology dictate whether a gene is cryptic or not.

To gain insight into the mechanism of formation of cryptic TSSs, we surveyed TF binding sites under unique PRO-cap peaks by performing an unsupervised motif analysis using HOMER^39^. The top 10 statistically significant motifs enriched at cryptic initiation sites were plotted as bubble plots (Fig. S4.1C-D). Specifically, we detected motifs bound by Activator Protein 1 (AP-1) family TFs such as ATF3 and BATF, and Erythroblast Transformation Specific (ETS) factor PU.1 (on an ETS/AP-1 composite motif) with very high significance in senescent cells (Fig. 5A). The AP-1 family of factors includes a cohort of basic leucine zipper proteins of the JUN (c-JUN, JUNB, JUND), FOS (c-FOS, FOSB, FRA1 and FRA2), MAF (c-MAF, MAFB, MAFA, MAFG/F/K and NRL) and ATF (ATF2, LRF1/ATF3, BATF, JDP1, JDP2) sub-families. AP-1 factors homo- or hetero- dimerize to bind DNA using the stretch of basic amino acids. c-JUN is the most potent transcription activator in the group^40^. We analyzed RNA-seq data to identify genes encoding AP-1 factors or their known interacting partners that increase in expression during senescence. The data revealed that expression of c-JUN, JUND, ATF3 and the group of small MAF proteins (MAFG/F/K) was elevated in senescence (Fig. 5B).

**Fig. 5:**
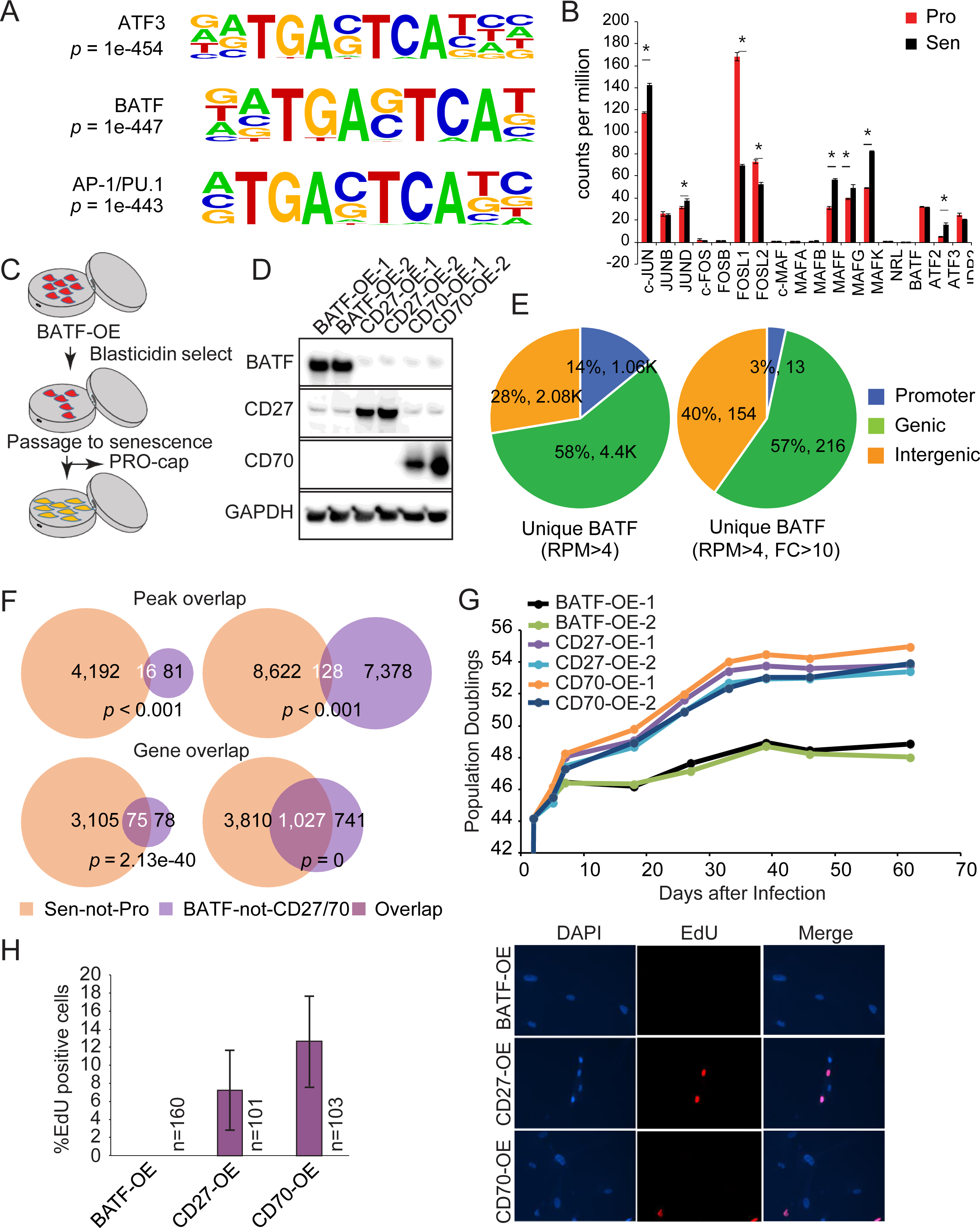
The AP-1 family of pioneer factors drive the formation of cryptic TSSs. **(A)** Top three most significant consensus motifs from HOMER motif analysis of a 300 bp region surrounding a cryptic TSS. *p-*values of enrichment are reported. **(B)** Expression analysis of AP-1 family of TFs and their interactors from two biological replicate RNA-seq datasets of proliferating and senescent cells^54^. Error bars represent standard deviation. * represents *p-*value < 0.05 in a two-tailed t-test. **(C)** Schematic showing experiment with BATF overexpression. **(D)** Immunoblot of whole cell lysates from IMR90 cells overexpressing BATF (AP-1 factor), CD27 (control) or CD70 (control). **(E)** Compartment analysis showing the genic enrichment of cryptic TSSs in cells overexpressing BATF. The left pie-chart shows BATF peaks with >4rpm with no overlap to any CD27 or CD70 peaks. The right pie chart shows BATF-induced peaks with >4rpm and >10-fold change compared to CD27 and CD70. **(F)** Overlaps of peaks (top) and genes (bottom) containing cryptic TSSs in senescence and in BATF overexpression. The *p-*values are from hypergeometric tests using all genes with PRO-cap peaks in 3’ half as universe. **(G)** Replicative lifespan curves of IMR90 cells overexpressing BATF, CD27 or CD70. **(H)** EdU incorporation assays of cells expressing BATF compared to those expressing CD27 or CD70 at endpoint. The quantification of EdU positive cells (from two independent biological replicates) for each sample is represented in the left bar graph. “n” represents the number of cells counted. Representative images are shown on the right.

To directly test whether AP-1 family pioneer factors can drive the senescence phenotype via cryptic transcription, we hypothesized that any AP-1 factor, if sufficiently expressed in senescence, would be recruited to AP-1 sites (including cryptic TSSs) to promote transcription. We thus overexpressed BATF, as an AP-1 family factor not expressed in IMR90 but with prominent roles in B and T cell function^41^, in PD 44 proliferating IMR90 cells (Fig. 5C-D) and performed PRO-cap. As controls, we overexpressed the membrane-bound ligand-receptor pair CD27 and CD70, known to be involved in B and T cell function but not nuclear in localization.

Cells overexpressing BATF generated new cryptic sites primarily within genes (Fig. 5E) as in proliferating and senescent cells. Further, these new cryptic sites largely overlapped with cryptic genes formed in senescence (Fig. 5F, lower), with the actual intragenic sites showing a smaller but still significant overlap (Fig. 5F, upper) possibly due to factors other than BATF also contributing to their formation. Importantly, the enhanced intragenic transcription in BATF overexpressing cells dramatically reduced growth compared to controls (Fig. 5G). This also compromised cellular health: at near-endpoint (assayed a few PDs before growth cessation of controls), there was no incorporation of EdU in the BATF overexpressing cells (Fig. 5H). Taken together, this data suggests that AP-1 factors, which are activated during the integrated stress response^42^, are able to drive cryptic transcription, and lead to defects in growth.

### Cryptic transcription is observed in aged female but not male murine livers

We next investigated whether cryptic transcription is also observed in aged tissues. We chose to analyze cryptic transcription in the liver because it is a relatively homogenous organ consisting of ∼70% hepatocytes and therefore unlikely to be grossly affected by cellular compositional differences during aging. We generated PRO-cap libraries from frozen livers of young (n = 4; age 2-4 months) and old (n = 4; age 22-25 months) mice including both genders. The run-on reaction was performed on isolated chromatin using a method called ChRO-cap^43^ (Fig. S5.1). By imposing similar filtering criteria as in senescent cells (intragenic peaks that are in the 3’ half of the gene and at least 3.5 Kb away from any other TSS), we detected numerous intragenic PRO-cap peaks in the old mice (Fig. 6A-B). For the ChRO-cap dataset however, we imposed an rpm cutoff of 10 rather than 4 because visual inspection of peaks revealed higher background from frozen tissues. With this cutoff, we identified “Young-not-Old” and “Old-not-Young” peaks unique to each condition. Of note, we detected far more unique cryptic TSSs in the old than we did in the young (Fig. 6B and Table S1). Example Gene Browser shots of intragenic cryptic TSSs in old mice are shown in Fig. 6C. Of particular interest, segregating the PRO-cap data by sex revealed a surprising phenomenon of cryptic initiation almost exclusively in female mice (Fig. 6D); while we do not understand the reason behind this bias, it may provide a molecular signature to assess sex differences in aging studies, as reported for calorie restriction^44^. Directional heatmaps revealed that most of the Old-not-Young cryptic TSSs generated unidirectional transcripts, suggesting that as in senescent cells, they reflect promoter, rather than enhancer transcription (Fig. 6E). As with proliferating and senescent cells, we found that cryptic sites in mice livers also occurred in long genes (Fig. 6F). We surveyed TF binding sites underlying the cryptic TSSs in old mice and found strong enrichments in motifs for Oct2/4 and the ETS family of TFs (Fig. 6G). One enriched motif was for the pioneer factor PU.1^45^, suggesting that pioneer factor mediated cryptic transcription during aging may be a conserved mechanism. A GO analysis of the genes harboring cryptic sites in old mouse livers revealed pathways connected to intracellular signal transduction, GTPase activity and immune function (Fig. S6.1B), similar categories to senescent human IMR90 cells. Given the limited number of Young-not-Old genes, only one significant GO category was detected (Fig. S6.1A).

**Fig. 6:**
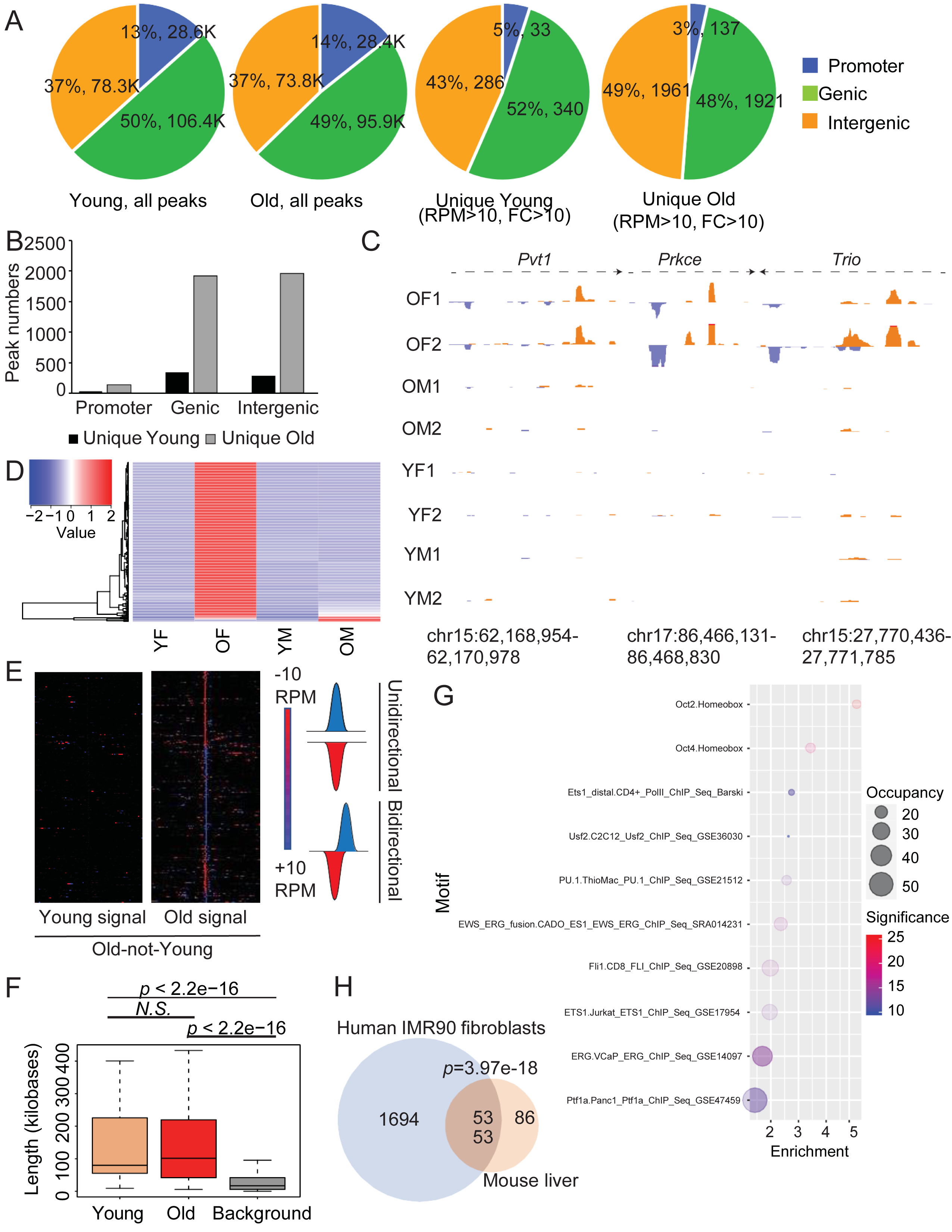
Aged mouse livers show evidence or intragenic cryptic transcription. **(A)** Compartment analysis of all (first two pie charts) and unique (last two pie charts) ChRO-cap peaks in proliferating and senescent cells as called by HOMER. Unique peaks were filtered to retain those that had a signal above 10rpm and a 10-fold enrichment by an AUC measurement. Promoters were assigned as 1 Kb around the TSS, genic as everything in the gene except the promoter region and intergenic as everything but promoter and gene. Proportions of peaks in a specific compartment are denoted as % and total numbers of peaks are indicated. **(B)** Peak numbers from the last two pie charts are plotted as a bar graph. **(C)** Browser track views of new ChRO-cap peaks detected in old mice livers. **(D)** Heatmap showing the ChRO-cap signal segregated by sex. **(E)** Heatmaps of genes that show evidence of cryptic transcription in young and old livers, centered around the unique ChRO-cap peaks detected in the old. Blue and red indicate signal in plus and minus strands respectively. **(F)** The length of the Young-not-Old and Old-not-Young cryptic genes is plotted in relation to all genes in the mouse genome. *p-*values are reported based on a Mann-Whitney U test. **(G)** Bubble plots of motif analysis (Known motifs from HOMER) of sequences under Old-not-Young cryptic peaks. The significance (-log (*p*-value)) is depicted as a color scale with warmer colors showing the greatest significance. The area of the circle is proportional to the % sites occupied. The fold-enrichment (x-axis) is the ratio of % sites occupied to % background sites occupied. **(H)** Venn diagram showing that 53 genes are orthologous between human and mouse. *p* value is from a hypergeometric test.

Since the GO terms observed in human senescence and aged mouse livers were similar, we investigated whether the cryptic genes in human senescent cells were orthologous to cryptic genes in aged mouse livers. Orthologs are defined as genes in different species that have evolved through speciation events only and thus are likely to perform similar biological functions. Interestingly, we observed that a significant proportion of cryptic genes (53 genes) in old mice were orthologous to genes showing upregulated cryptic in human senescent cells (Fig. 6H). A GO analysis of age-related cryptic genes showed that the orthologous genes are related to signal transduction (Fig. S6.1C).

## Discussion

In this study, we employ base-pair resolution run-on methodologies^25^ to directly probe for aberrant transcription initiation events in senescent human fibroblasts and aged murine livers. Our results reveal that as in model organisms such as yeast and worms^3^, spurious initiation from within gene bodies is a frequent event during mammalian aging. To understand the cause of spurious initiation, we performed mechanistic studies in human senescent fibroblasts. We profiled the chromatin architecture at these cryptic sites and found evidence of “fuzzy” TSS-like structures forming primarily over poised intronic enhancers (Figs. 1-3). Our results thus indicate that an enhancer-to-promoter conversion occurs in senescence and is driven by pioneer factors that then enables deposition of H3K4me3, p300/CBP recruitment and H3K27ac (Figs. 2 and 3). Upon deletion of their constitutive promoters, some genes are capable of transcribing multi-exonic RNAs called meRNAs, by using their intronic enhancers as alternative promoters^46^ suggesting that enhancers have the potential to act as promoters.

In senescent human fibroblast cells, the AP-1 family of pioneer factors emerged as potential drivers of the cryptic phenotype. We show that AP-1-driven cryptic transcription causes cells to undergo a stress-induced growth arrest phenotype (Fig. 5). We also discover ETS family of TFs as potential drivers of cryptic phenotype in both senescent cells (Fig. S4.1D) and old mouse livers (Fig. 6G). ETS factors such as PU.1 have been shown to cooperate with AP-1 factors by binding to composite motifs and forming ternary complexes^47,48^.

Recently, the role of widespread and ubiquitously-expressed AP-1 and ETS family TFs in enhancer and chromatin looping have been proposed to drive reprogramming and cancer^49^. Our study suggests these factors may cause changes in the genic landscape and affect fidelity of transcription initiation; the cause of their increased activity in senescence and aging remains to be investigated. One hypothesis is upregulation of the integrated stress response which triggers binding of heterodimeric complexes of ATF4 and c-JUN to AP-1 response elements^42^. In support of this study, two very recent reports have indicated the role of AP-1 TFs in enhancer remodeling and chromatin accessibility in senescence^50,51^.

Genes prone to age-related cryptic transcription tend to be long, actively expressed, and have large intronic sequences (Fig. S1.2). Cryptic TSSs tend to form over these large introns in areas of low DNA and histone methylation that are otherwise thought to provide protection against inappropriate initiation^3,4,52^ (Figs. 4 and S3.2). However, in general, these genes have high levels of DNA and histone methylation (Fig. 4), perhaps having evolved to protect them from aberrantly initiating transcription from non-canonical TSSs or simply epiphenomenal to transcriptional activity. The genes encode for proteins involved in signal transduction pathways that contribute to SASP and cancer, reiterating the importance of maintaining their proper transcription fidelity (Fig. S4.1). The recapitulation of cryptic transcription events in aged mouse livers and in orthologous genes (Figs. 6 and S6.1) imply that loss of the transcription fidelity is a conserved feature of aging cells. Targeting the pathways that contribute to this molecular phenotype can be used to develop therapies against age-associated structural and functional damage in tissues.

## Online Methods

### Experimental methods

#### Cell culture

IMR90 cells, obtained from Coriell Institute for Medical Research, Camden, NJ, were grown in standard tissue culture medium (DMEM with 10% FBS and 1% penicillin/streptomycin; Invitrogen) at 3% oxygen. Senescence was induced by replicative exhaustion. Briefly, for each passage, cells were washed with PBS, trypsinized at 37 °C for 5 minutes, and plated on fresh 10 cm plates at 0.5-1e6/plate or fresh 15cm plates at 1-2e6/plate.

Cells were counted with a Countess automated cell counter (Life Technologies), and the numbers were recorded for growth curve generation. Cell viability record was maintained via Trypan Blue staining. Cells were maintained in dishes for 2 weeks to ensure growth termination/senescence at endpoint. Senescence-associated β-gal expression was tested to confirm senescence using a kit (EMD Millipore). For overexpression experiments, IMR90 cells were infected with lentiviral constructs designed against selected targets in medium containing 8ug/ml polybrene. After 24 hours, the cells were replaced with polybrene-free media and after another 24 hours, selection was initiated with addition of blasticidin. After selection was complete, the media was changed back to standard medium.

#### Animals

C57/BL6J young mice were ordered from Jackson Laboratories, old mice were obtained from the National Institute on Aging rodent colony. Mice were fed ad libitum on a standard chow diet and handled following institutional regulations and guidelines. Both sexes were included in the study.

#### Quantitative PCR (qPCR)

To validate PRO-cap peaks, qPCR was performed using 7ng of PRO-cap library against genomic DNA standards (100ng – 10e-4ng) qPCR with FastSYBR (Applied Biosystems).

#### Western blotting

Cells were lysed in buffer containing 50 mM Tris pH 7.5, 0.5 mM EDTA, 150 mM NaCl, 1% NP40, 1% SDS, supplemented with 1X Halt Protease inhibitor cocktail (Thermo Scientific). The lysates were briefly sonicated, and cleared by centrifugation at max speed for 10 minutes at 4°C. The supernatants were quantified using the BCA kit (Pierce) and subjected to electrophoresis using NuPAGE 12% Bis-Tris gels. The proteins were transferred to a 0.2-micron nitrocellulose membrane using a wet transfer apparatus (BioRad) for 1.5 hours at 100V (4°C). 5% milk in TBS supplemented with 0.1% Tween 20 (TBST) was used to block the membrane at room temperature (∼25 °C) for 1 h. Primary antibodies were diluted in 3% BSA in TBST and incubated at 4 °C overnight. The list of antibodies used in this study is listed in Table S3. The membrane was washed three times with TBST, each for 10 minutes, followed by incubation with HRP-conjugated secondary antibodies (BioRad) at room temperature for 1 h, in TBST. The membrane was washed again three times and imaged by a Fujifilm LAS-4000 imager.

#### EdU assay

EdU assays were performed using the Click-iT EdU Alexa Fluor 488 Imaging Kit (Thermo Scientific) to measure the percentage of cycling cells. Briefly, 2e5 cells were plated on poly-L-lysine (Sigma) coated coverslips in a 24-well plate and labeled with 1X EdU solution overnight (24 hours). The cells were fixed with 4% PFA and permeabilized using 0.5% Triton X-100 in PBS. EdU labeled cells are then detected by a copper- catalyzed ‘‘click’’ reaction with a small molecule conjugated to Alexa Fluor followed by DAPI staining. The coverslips were mounted with ProLong Gold Antifade reagent (Invitrogen) and EdU incorporation calculated as a percentage of DAPI-stained cells also showing Alexa Fluor signal imaged by a fluorescent microscope (Nikon Eclipse 80i).

#### Precision Run-On-sequencing (PRO-seq and PRO-cap) and Chromatin Run-On sequencing (ChRO-seq)

PRO-seq/cap was performed on proliferating and senescent IMR90 cells as described previously^25^. Briefly, ∼12e6 trypsinized cells were permeabilized and stored in storage buffer (10mM Tris-HCl pH 8.0, 25% glycerol, 5mM MgCl2, 0.1mM EDTA, 5mM DTT) at -80°C until use. The permeabilized cells were then resuspended in 1X nuclear run-on reaction mix containing biotinylated rCTP, rATP, rGTP and rUTP for 3 minutes at 37°C. RNA was extracted, fragmented and subjected to biotin enrichment. The enriched nascent RNA population was then ligated to a 3’ adapter sequence and subjected to a second round of biotin enrichment. The 5’ ends of the RNA were decapped, phosphorylated, ligated to 5’ adapter and subjected to a third biotin enrichment. The RNA was then reverse transcribed and PCR amplified with appropriate index primers. The libraries were purified by extraction from an 8% polyacrylamide gel, quantified by Bioanalyzer (Agilent) and qPCR (Kapa Biosystems).

The PRO-seq/cap run was performed on the NextSeq 500 platform (Illumina). ChRO-seq was performed as described previously^43^ with ∼250mg frozen liver tissue from young and old mice.

#### Chromatin immunoprecipitation-sequencing (ChIP-seq)

ChIP was performed as described previously from ∼12e6 cells^17^. Briefly, cross-linked cells were lysed, and the chromatin was sheared to an average size of <500 bp using a Covaris S220 Ultrasonicator. 2ug of sonicated chromatin was used per immunoprecipitation reaction (see Table S3 for a list of antibodies), and 10% of the amount was saved as input. Immunoprecipitation was performed using protein A Dynabeads (Life Technologies). For ChIP-seq, 1ng of DNA from immunoprecipitation and input was used to prepare libraries with the NEBNext Ultra II kit (New England Biolabs). The libraries were checked for quality and quantity by Bioanalyzer (Agilent) and qPCR (Kapa Biosystems) respectively. Multiplexed libraries were sequenced on a single lane of the NextSeq 500 platform (Illumina).

#### Assay for Transposase Accessible Chromatin (ATAC-seq)

ATAC-seq was performed using a protocol from the Greenleaf Lab^53^. 100,000 proliferating or senescent IMR90 cells were used to generate libraries in two biological replicate experiments.

#### Whole genome bisulfite sequencing (WGBS)

WGBS from proliferating and senescent cells has been previously described^54^. The methylation data is available under GEO accession number GSE48580.

### Computational Methods

#### PRO-seq/cap analysis

PRO-seq/cap data were aligned to the NCBI v37 assembly of the human genome (hg19) using ProSeqMapper (Danko lab, https://github.com/Danko-Lab/tutorials/blob/master/PRO-seq.md) with default parameters (no UMI barcode filter) and a BWA index. To call PRO-cap peaks, we used HOMER’s findPeaks method with -style tss on output BAM files with no filtering on alignment scores. To obtain a stringent list of new PRO-cap peaks in the senescence condition, this initial list was further filtered to retain only those peaks that have a signal >4rpm (HOMER normalized tag counts) for human or >10rpm for mouse and >10-fold enrichment in an area-under-the-curve analysis. A pseudocount of 0.01 was added for fold change calculation to avoid division by zero. Any peak from this list that intersected (BedTools) with a peak from the other condition was removed. Finally, all peaks within 3.5kb of the nearest TSS (RefSeq or GENCODE) and peaks which did not fall in the 3’ half of a gene body were also removed. The 3.5kb distance cutoff was chosen by assessing the distribution of distances of all ENCODE Broad chromHMM elements^27^ in state 1 or 2 (promoters) to the nearest RefSeq TSS; roughly 2/3 of elements are within 3.5kb.

### Compartment Analysis

We defined promoters as a 1kb region upstream of the TSS. The gene region is all gene bodies without any promoters. Intergenic is defined as any genomic space neither gene nor promoter. Overlaps were determined using BEDtools intersect.

### Assessment of bidirectionality

PRO-cap enrichment was scored for a 5kb window in 50bp increments centered on all cryptic sites, measuring + and – reference strand-aligned tags separately. Clustering was performed by first taking the sum of the + and – strand standardized scores for the central 300bp region, then clustering on these two columns in R using hclust and dividing into 10 clusters using cutree. Heatmaps represent non-standardized scores sorted vertically using this clustering, scaling the maximum enrichment at 10rpm. To determine bidirectionality, we scored a 300bp window around each cryptic site for PRO-cap + and – strand aligned tag enrichment and computed a score max(log10(+/–), log10 (–/+)), with larger scores indicating unidirectional transcription and lower scores indicating bidirectional transcription. Data were plotted using ggplot2’s geom_density() function and the distribution difference between Pro-not-Sen and Sen-not-Pro sites was assessed for significance using a permutation test (coin package, independence_test() function).

### ATAC-seq analysis

ATAC-seq alignment was performed using bowtie2 on hg19 keeping only uniquely aligned tags (-k 1) with 1 mismatch in the seed (-N 1). SAM files were filtered by discarding tags where a segment was unmapped (-F 4) and those failing a MAPQ filter of 5 (-q 5). Replicate ATAC-seq samples were pooled after alignment and tracks were made after randomly downsampling the proliferating data set to match the coverage of the senescent data set.

### ChIP-seq analysis

ChIP-seq data were processed as described previously^55^. Briefly, data were aligned to human genome assembly GRCh37/hg19 using bowtie2 (v2.1.0) with parameters --phred33, -N 1. Data were sorted and filtered using samtools (v0.1.19) to remove poor alignments (-F 4 -q 10) and PCR duplicates were removed with samtools rmdup. BEDs were created from BAM files and reads aligning to the mitochondrial chromosome were removed using grep while reads falling into ENCODE blacklist regions were removed using BEDtools intersect -v. Finally, for the Shah et al data set^17^, H3 background control samples were randomly downsampled to match the coverage of the H3K4me3 chIP.

### AUC

The Area Under the Curve (AUC) for a given region in a given sample is defined as the number of tags aligned to that region multiplied by the rpm of the data set and 1000, divided by the length of the region. For ChIP-seq data, a similarly adjusted input score is subtracted from the ChIP.

### Boxplots, Heatmaps and Metaplots

All boxplots were created using R’s boxplot() function. PRO-cap heatmaps are plotted using a python script reliant on the Python Imaging Library (pillow module); vectors spanning 5kb windows with 100 50bp increments are scored by calculating the stranded AUC value in each increment for both conditions (proliferating and senescent). All metaplots were created by taking non-stranded AUC vector output of all locations and averaging for each increment for proliferating and senescent separately. For methylation meta- plots, genes were divided into 100 equal-sized bins and scored similarly; DNA methylation is the average of % methylated per bin, rather than AUC. The heatmap of Old-not-Young cryptic sites in mouse was created using a 300bp window and standardized in R, plotted with heatmap.2() from the gplots library.

### WGBS data analysis

Bisulfite sequencing reads were assessed for quality using FastQC (version 0.10.0) prior to having all adapters and low-quality sequence tails removed using trim-galore (version 0.3.0).

Sequenced reads were then transformed in silico to fully bisulfite-converted forward (C→T) and reverse (G→A) reads. The converted sequences were aligned against a converted UCSC (hg19) genome in each combination: (1) forward (C→T) reads align to forward (C→T) genome; (2) reverse (G→A) reads align to reverse (G→A) genome; (3) forward (C→T) reads align to reverse (G→A) genome; (4) reverse (G→A) reads align to forward (C→T) genome. During the library preparation process, genomic fragments representing alignments (3) and (4) are generated in the PCR step; however, they are not sequenced and only fragments corresponding to alignments (1) and (2) are retained. Alignment was carried out using Bismark [2] (version 0.10.1), based on the Bowtie2 [1] aligner (version 2.1.0) on the UCSC hg19 human genome.

For each aligned sequence tag, the original unconverted sequence was compared against the original unconverted reference genome and the methylation status was inferred. Sequences aligned from (1) and (2) gave information on cytosines on the forward and reverse strands respectively.

To remove PCR bias, a deduplication step removed potential duplicate reads, where both ends of the fragment align to the same genomic positions on the same strand, only one of these reads was retained. To control for potential incomplete bisulfite treatment, any reads with more than three methylated cytosines in non-CpG contexts were discarded.

Processed reads were aggregated on a per CpG basis (number of bases read supporting methylated/unmethylated status). At this stage we collapsed CpG dyads using a bespoke script that combined methylated and unmethylated coverage scores for each CpG dyad into a single score for the cytosine on the forward strand, thus artificially increasing the coverage with a slight loss in spatial resolution. A two-tailed Fisher exact test was used to identify CpGs that were differentially methylated. Only CpGs with at least ten reads within each comparable condition were considered for testing. *p*-values were corrected using the Benjimini-Hochberg (BH-) FDR function to control false positives at a rate of 5%.

The percentage CpG methylation for any given window was calculated as the total number of cytosines sequence (at CpG sites for that window), divided by the total number of methylated and unmethylated cytosines (at CpG sites for that window), multiplied by 100.

### Motif

All motif analysis was done using HOMER (v4.6), with default background, size parameter set to 300bp and masking. Motif enrichment is represented using bubble plots (ggplot2 library, geom_point() function), taking the first 10 motifs from the known results and sorting by the enrichment score (% foreground : % background ratio). Significance (heat) is -log10(p-value) and occupancy is % foreground.

### RNA-seq analysis

RNA-seq datasets from the Adams lab^56^ were first trimmed using Trimmomatic-0.32 with parameters SE, LEADING:3, TRAILING:3, SLIDINGWINDOW:3:3 and MINLEN:30. The tags were single end aligned using STAR-2.3.0e with parameter alignMatesGapMax set to 2000. The aligned tags were filtered on a quality score of at least 10. The expression data was obtained using featurecounts-1.5.0 with parameters M, s0 over RefSeq genes.

### Gene Ontology analysis

Functional characterization of gene sets was performed using DAVID Bioinformatics Resources (http://david.abcc.ncifcrf.gov/home.jsp). Only GO terms with FDR<10 were taken into consideration. GO terms are represented using bubble plots made with ggplot2 (geom_point() function), taking the top 20 terms by FDR and sorting by fold enrichment.

### Orthology analysis

MGI’s vertebrate homology flat file (based on HomoloGene) was used to assign orthologs between human and mouse. Overlap significance was assessed using the hypergeometric calculation (phyper() in R) against the universe of all orthologous genes.

### Data accessibility

The GEO accession number for all genome-wide datasets generated in this article is GSE156829.

## Acknowledgments

We wish to thank Katherine Alexander and Digbijay Mahat for PRO-seq/PRO-cap/ChRO-cap guidance and members of the Berger lab for critical reading of the manuscript. We also want to thank Deepak Jha for insightful discussions. This work was supported by National Institutes of Health (NIH)/National Institute on Aging (NIA) grant P01AG031862 to S.L.B., American Heart Association grant 15POST21230000, AFAR Irene Diamond Transition Award DIAMOND 17113 and NIA IRP grant 1-ZIA-AG-000679-02 to P.S.

## Author Contributions

P.S. and S.L.B. conceptualized the work. P.S. generated most genome-wide datasets used in this study (except those already published as indicated in Table S2). C.L. performed functional experiments such as qPCR detection of PRO-cap peaks and replicative senescence assays with cells overexpressing BATF. G.E. performed PRO-cap on cells overexpressing BATF. G.D., E.K. and Y.L. performed bioinformatics analyses of data. N.R. processed WGBS data which was generated in the lab of P.D.A. D.C. S. provided concentrated lentiviral preparations used in this study. P.P.S. performed ATAC-seq and participated in discussions.

## Supplementary Materials

**Fig. S1.1:**
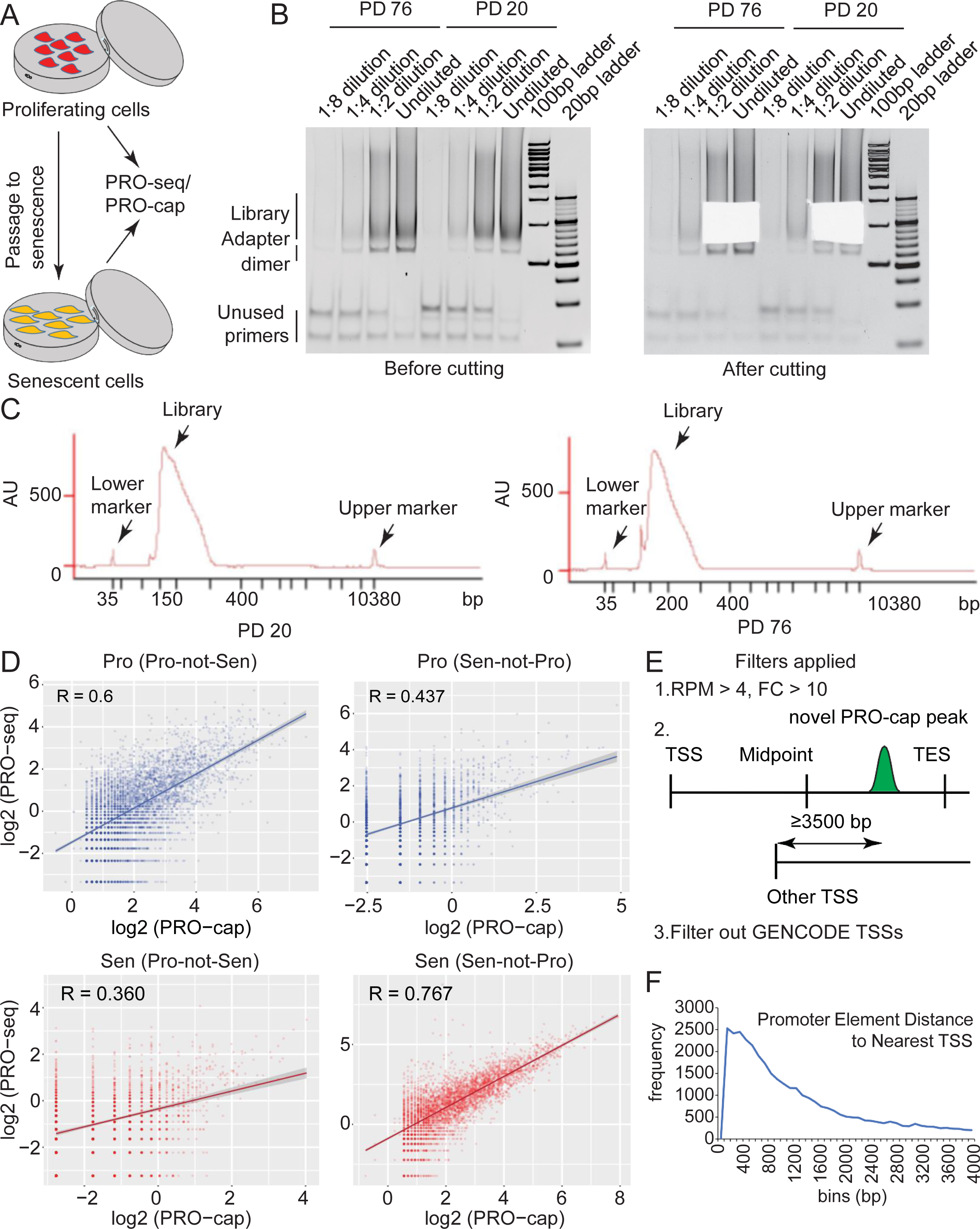
Construction of PRO-cap libraries from proliferating and senescent cells. **(A)** Schematic illustrating samples used for generating PRO-seq and PRO-cap libraries. **(B)** (Left) Representative image of 8% native polyacrylamide gels stained with ethidium bromide used for size selecting PRO-cap libraries from (A). The sizes of the library, adapter dimer and PCR primers are indicated along with size markers. (Right) Same as (B) except that the image was taken after cutting out the region containing the PRO-cap library. **(C)** (Left) Representative Bio-analyzer electropherogram of PRO-cap library from proliferating cells. (Middle) Same as (Left) except the library is from senescent cells. **(D)** Scatter plots showing correlation of PRO-cap and PRO-seq peaks, the correlation coefficient R is indicated. **(E)** Filters applied to PRO-cap peaks to (1) select unique peaks in one condition, (2) select only gene-internal sites away from canonical promoters and (3) ensure non-coding start sites are not selected.

**Fig. S1.2:**
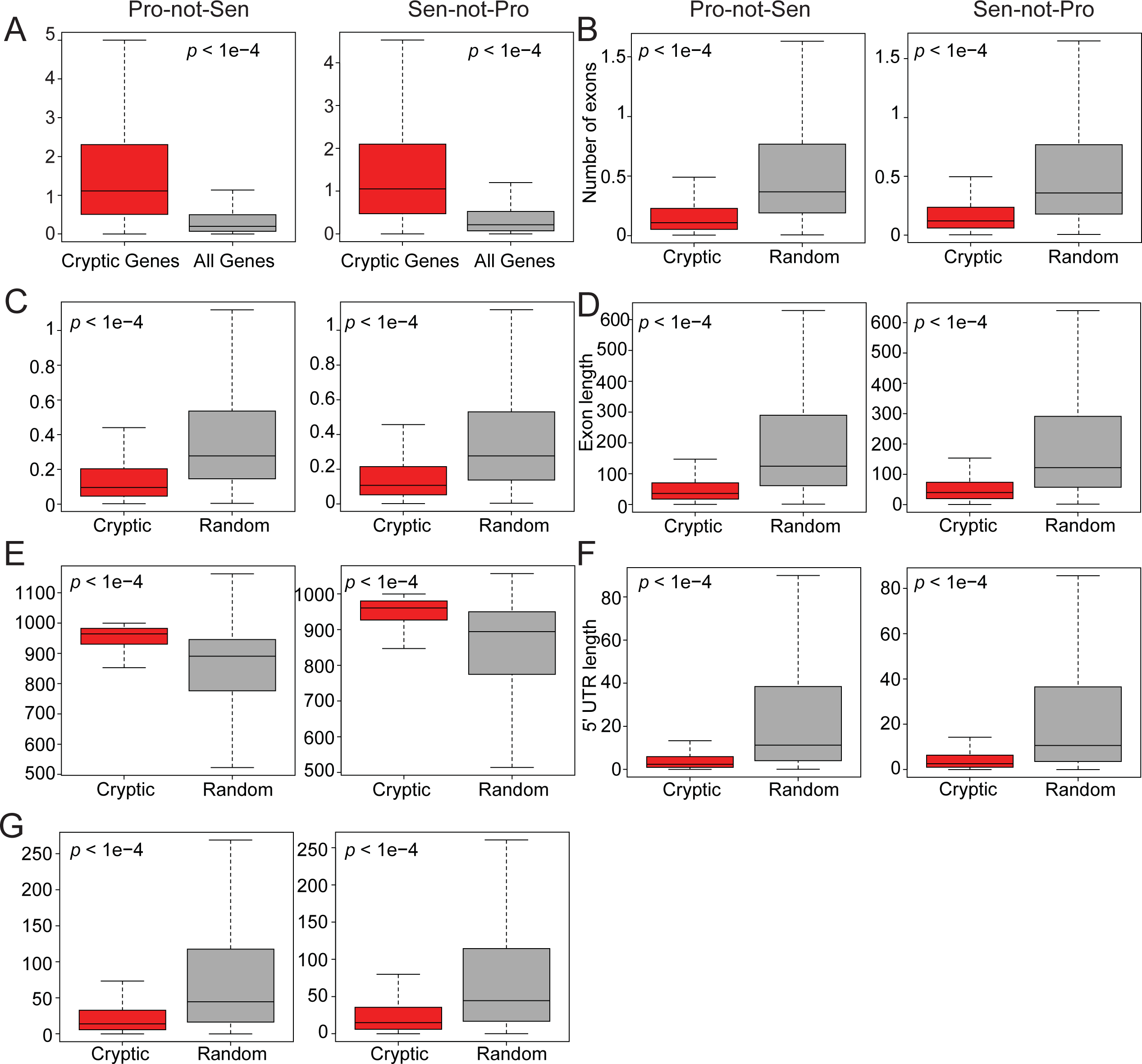
Features of cryptic genes. **(A)** The length of the cryptic genes is plotted in relation to all genes in the human genome. **(B-G)** The number of exons **(B)**, number of introns **(C)**, normalized length of exons **(D)**, normalized length of introns **(E)**, normalized length of 5’ UTRs **(F)** and normalized length of 3’UTRs **(G)** in cryptic genes is compared to an equal number of random genes. *p-*value is reported based on a coin library permutation test.

**Fig. S2.1:**
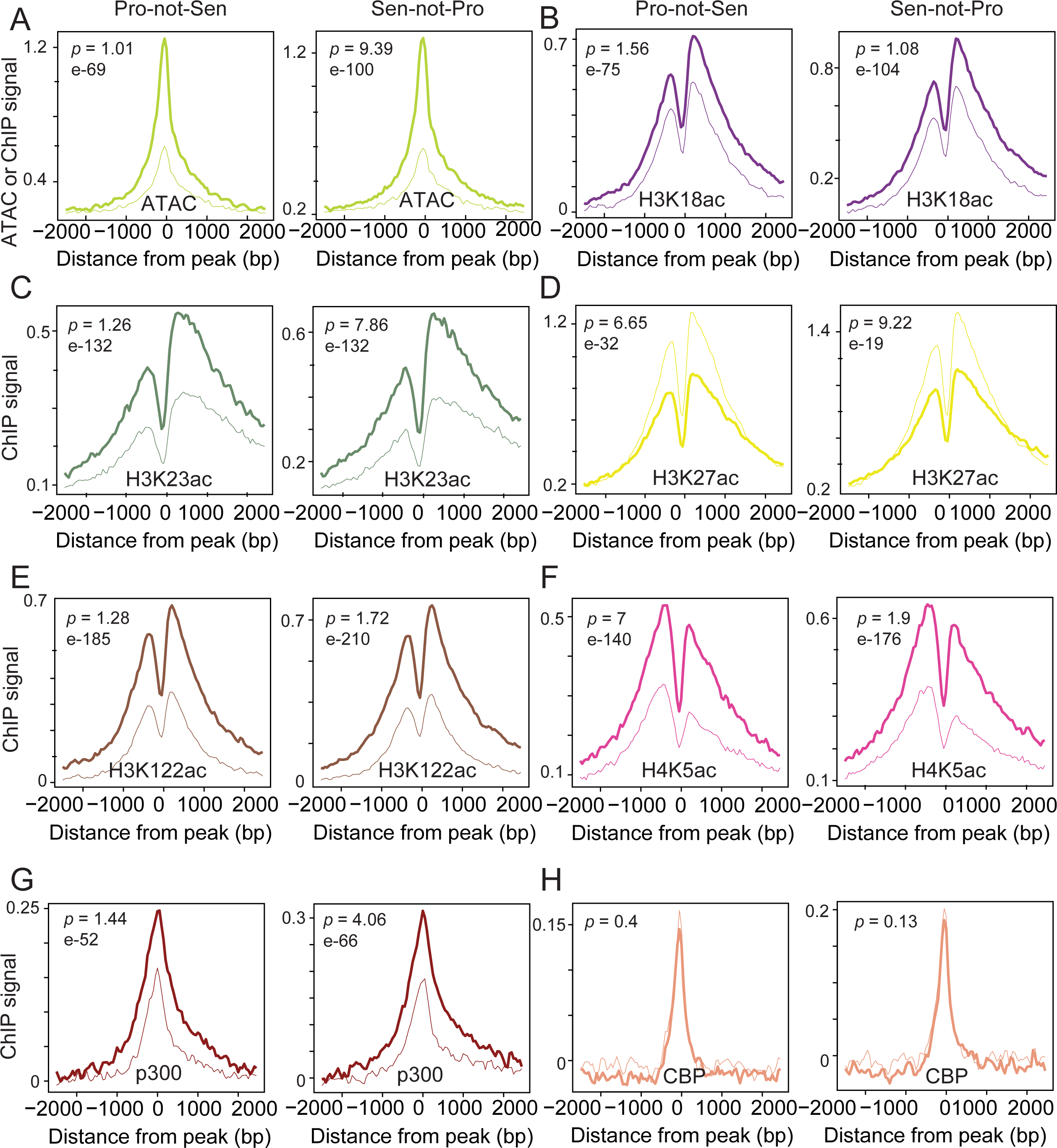
Chromatin acetylation profiles around canonical TSSs Metaplots of ATAC-seq. **(A)**, H3K18ac **(B)**, H3K23ac **(C)**, H3K27ac **(D)**, H3K122ac **(E)**, H4K5ac **(F)**, p300 **(G)**, and CBP **(H)** signal are shown in a 2 Kb region surrounding a canonical TSSs. Thin lines represent proliferating cells and thick lines senescent cells. *p-*values are reported based on a Mann-Whitney U test. Two outlier regions (chr2:88122786-88127786 in Pro-not-Sen and chr5:79947966-79952966 in Sen-not-Pro) with very high signal was omitted in (A).

**Fig. S3.1:**
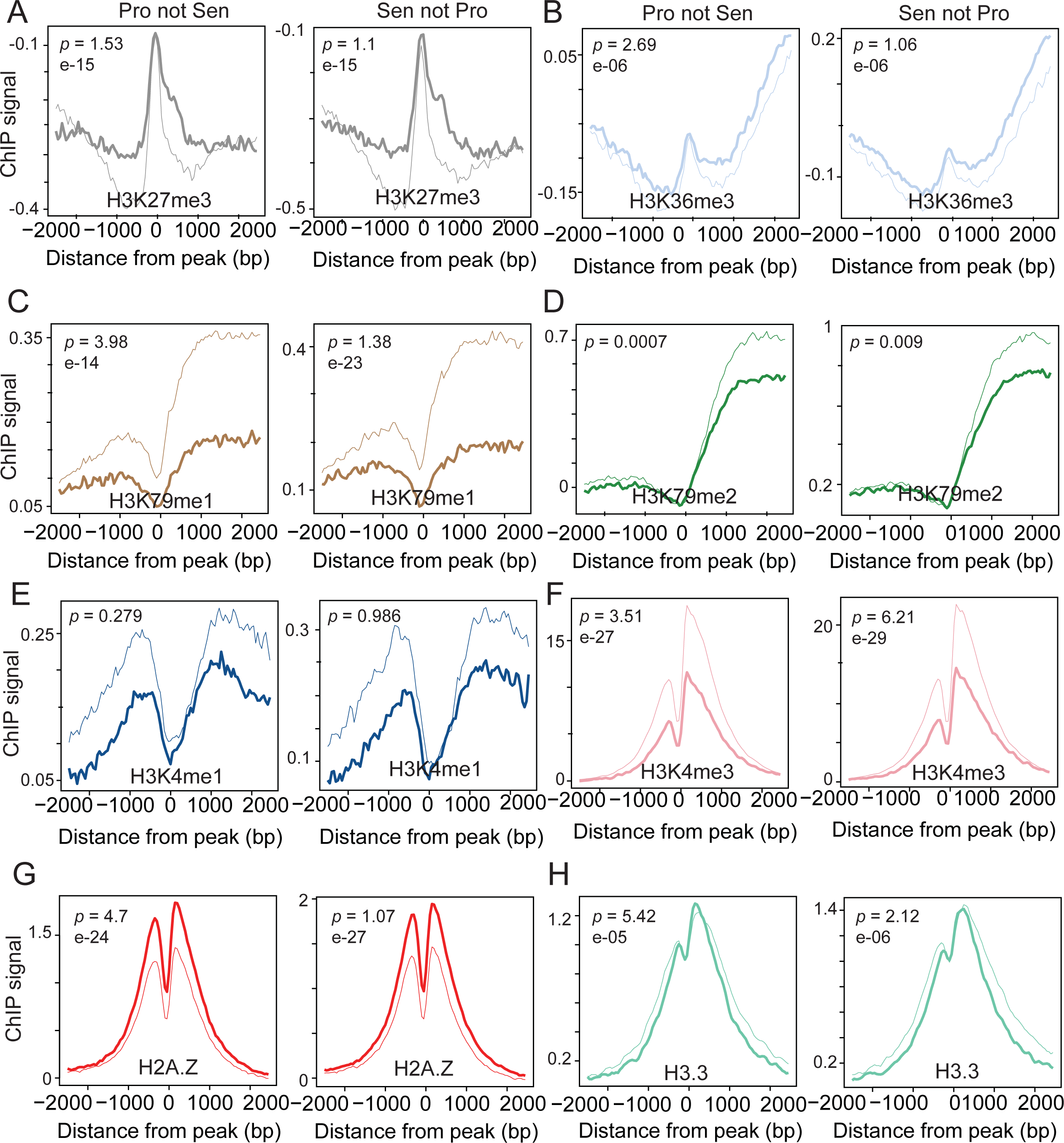
Chromatin methylation profiles around canonical TSSs Metaplots of H3K27me3. **(A)**, H3K36me3 **(B)**, H3K79me1 **(C)**, H3K79me2 **(D)**, H3K4me1 **(E)**, H3K4me3 **(F)**, H2A.Z **(G)**, and H3.3 **(H)** signal are shown in a 2 Kb region surrounding canonical TSSs. Thin lines represent proliferating cells and thick lines senescent cells. *p-*values are reported based on a Mann-Whitney U test.

**Fig. S3.2:**
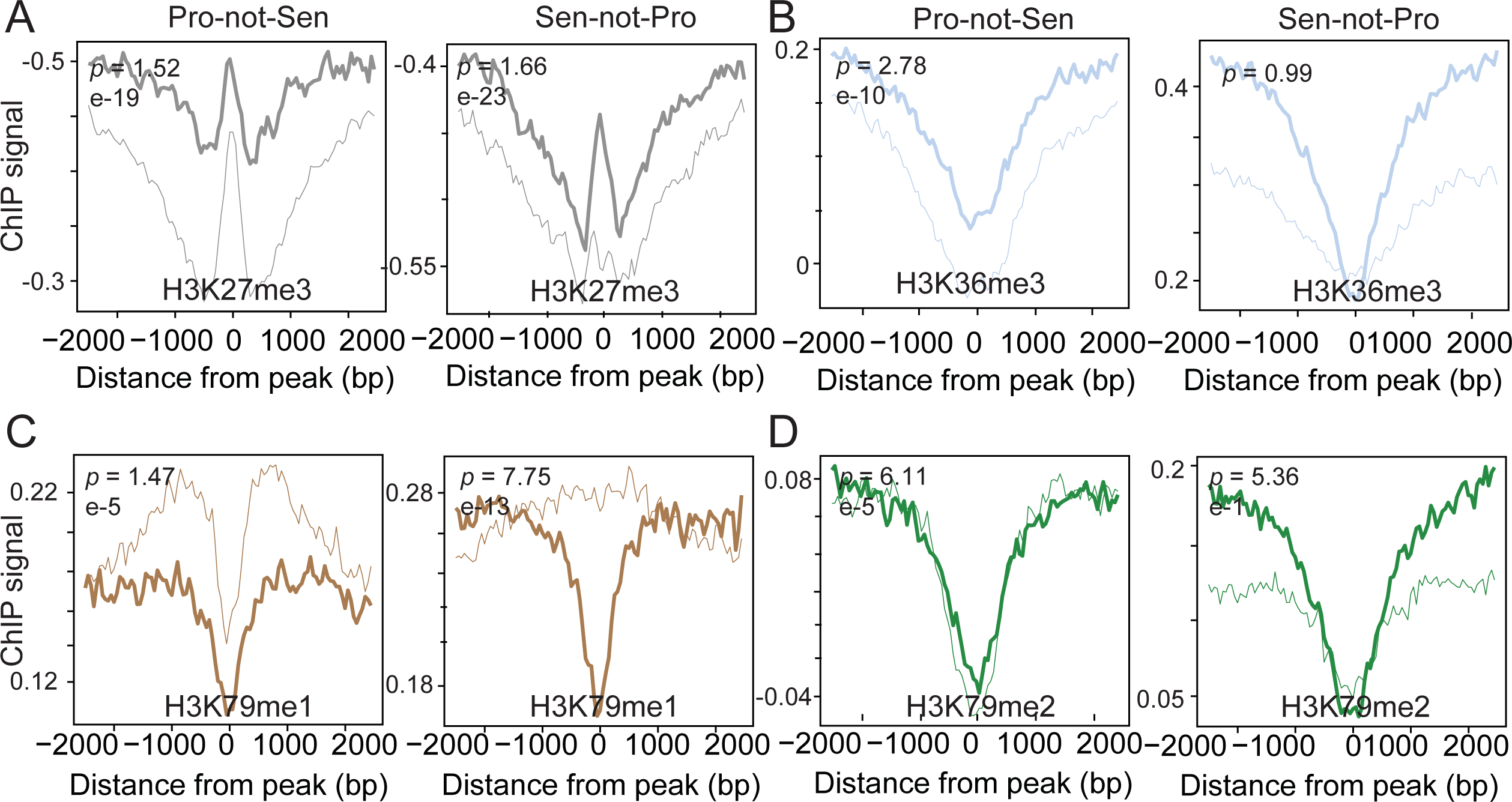
Cryptic TSSs show low histone methylation signal Metaplots of H3K27me3. **(A)**, H3K36me3 **(B)**, H3K79me1 **(C)**, and H3K79me2 **(D)** signal are shown in a 2 Kb region surrounding cryptic TSSs. Thin lines represent proliferating cells and thick lines senescent cells. *p-* values are reported based on a Mann-Whitney U test.

**Fig. S3.3:**
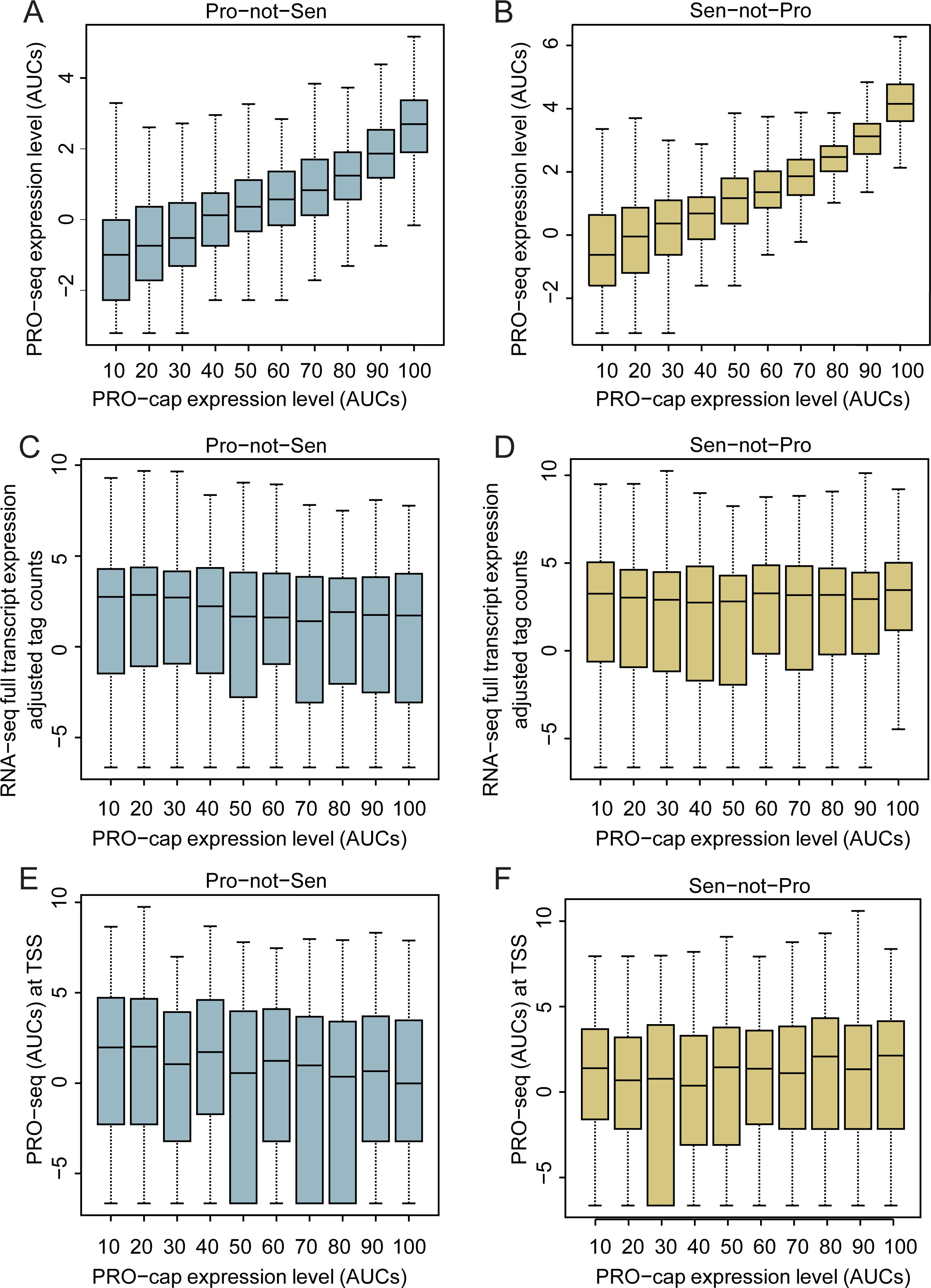
Cryptic TSSs are formed in actively transcribed genes. **(A-B)** PRO-cap signal at Pro-not-Sen TSSs **(A)** or Sen-not-Pro TSSs **(B)** were divided into deciles and the corresponding PRO-seq signal at the cryptic sites plotted. **(C-D)** Steady state RNA-seq signal in Pro-not-Sen **(C)** or Sen-not-Pro **(D)** cryptic genes was plotted across the deciles in A-B. **(E-F)** Same as C-D except PRO-seq signal at canonical TSSs of Pro-not-Sen cryptic genes **(E)** and Sen-not-Pro cryptic genes **(F)** is plotted. The decile-based *p-*values from a Mann-Whitney test are reported in Table S4. “Percentile Cutoff” indicates comparison of the bottom N% to the top 100-N%.

**Fig. S4.1:**
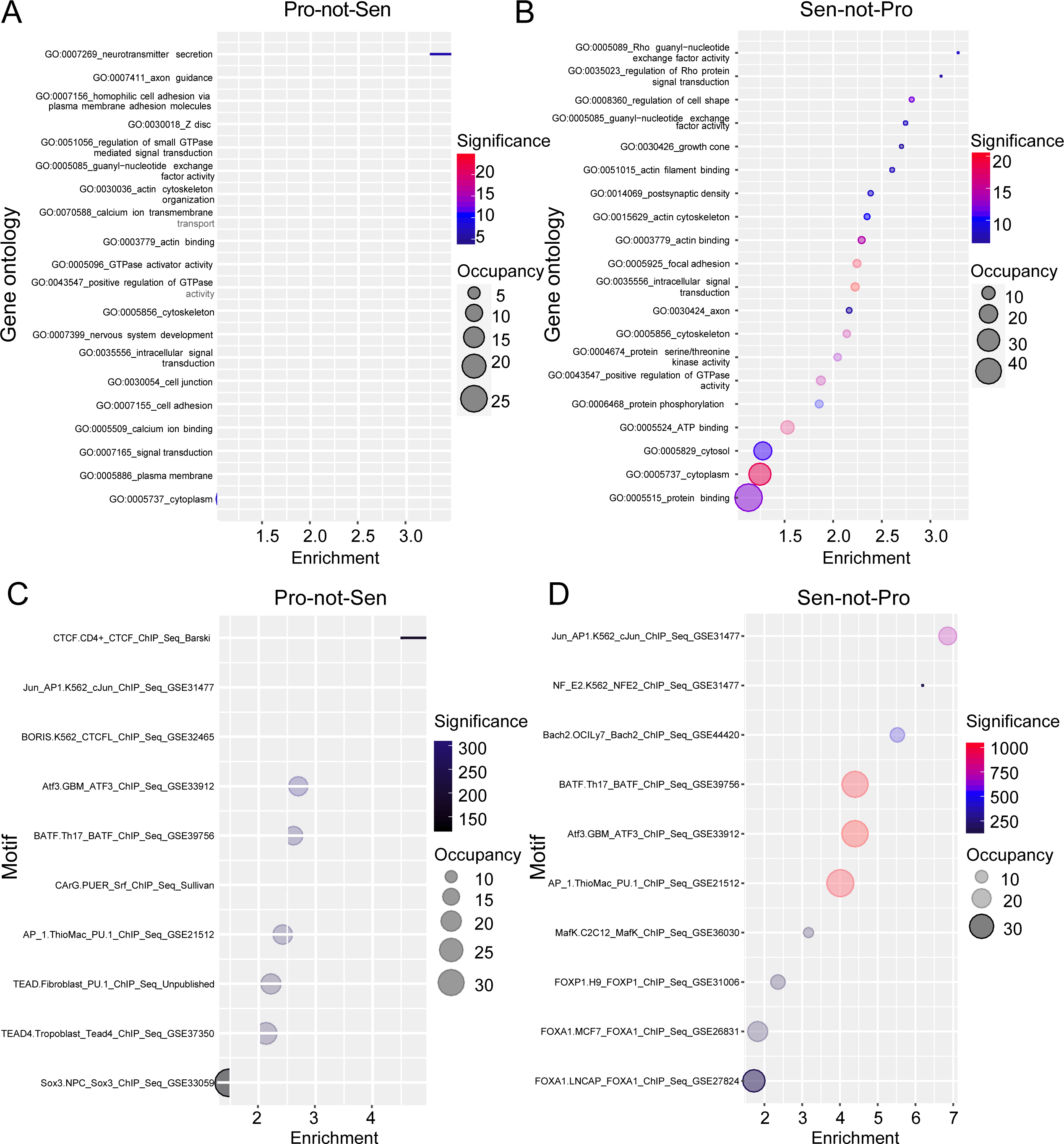
Gene Ontology and motif analysis of cryptic TSSs. **(A-B)** Gene Ontology analysis plotted as bubble plots of Pro-not-Sen **(A)** and Sen-not-Pro **(B)** cryptic sites. **(C-D)** Motif analysis (Known motifs from HOMER) of sequences under Pro-not-Sen **(C)** and Sen-not-Pro **(D)** cryptic TSSs. The significance (-log (*p*-value)) is depicted as a color scale with warmer colors showing the greatest significance. The area of the circle is proportional to the % sites occupied. The fold-enrichment (x-axis) is the ratio of % sites occupied to % background sites occupied.

**Fig S5.1:**
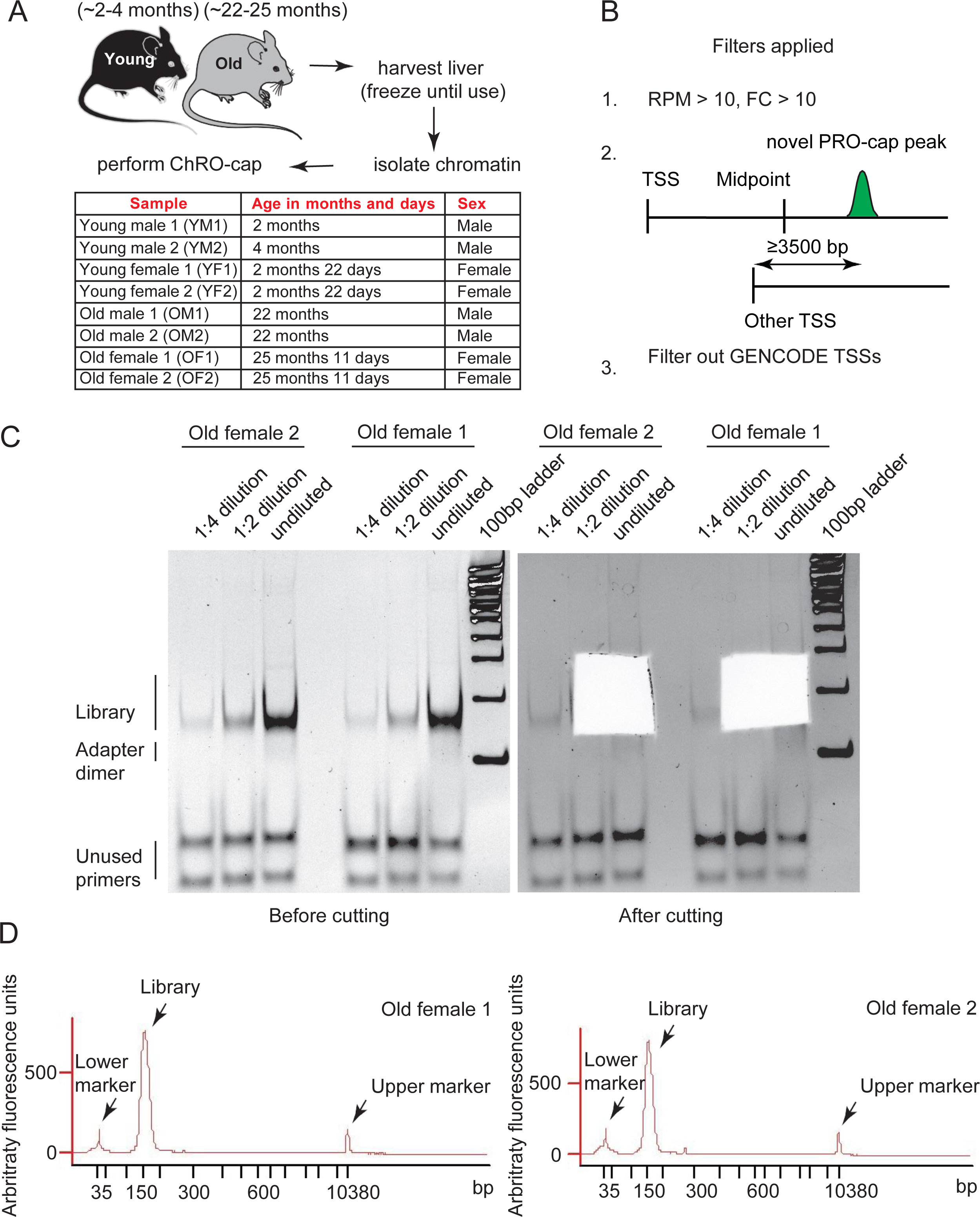
Construction of ChRO-cap libraries from young and old mice livers. **(A)** Schematic illustrating ChRO-cap strategy adopted for frozen mouse livers. Table shows the age and sex distribution of animals used for ChRO-cap. **(B)** Filters applied to ChRO-cap peaks to (1) select unique peaks in one condition, (2) select only gene-internal sites away from canonical promoters and (3) ensure non-coding start sites are not selected. **(C)** (Left) Representative image of 8% native polyacrylamide gels stained with ethidium bromide used for size selecting ChRO-cap libraries generated from old female livers. The sizes of the library, adapter dimer and PCR primers are indicated along with size markers. (Right) Same as (Left) except that the image was taken after cutting out the region containing the ChRO-cap library. **(D)** Representative Bio- analyzer electropherogram of ChRO-cap library from old female livers.

**Fig. S6.1:**
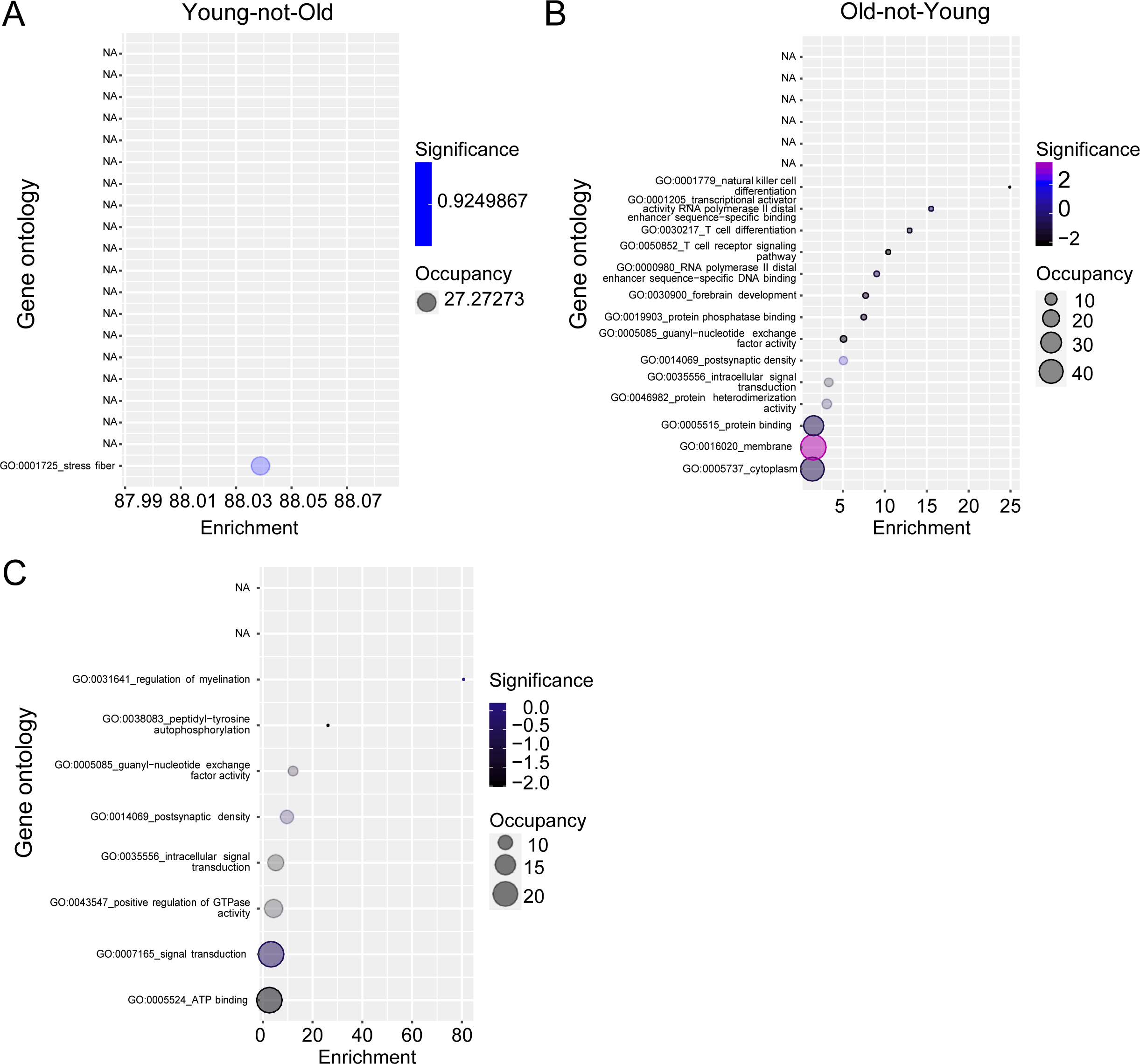
Orthologous genes in human and mouse upregulate cryptic transcription. **(A-B)** Gene Ontology analysis plotted as bubble plots of Young-not-Old **(A)** and Old-not-Young **(B)** genes that harbor cryptic TSSs. **(C)** Gene Ontology analysis plotted as a bubble plot for the 53 orthologous genes in Fig. 7H.

Table S1: List of Pro-not-Sen, Sen-not-Pro, Young-not-Old and Old-not-Young peaks. Table S1 is an excel file that accompanies this manuscript.

**Table S2:** Genome-wide data sets used in this study

| Datasets | Reference |
| --- | --- |
| PRO-seq | 55 |
| PRO-cap | This study |
| RNA-seq | 54 |
| ATAC-seq | This study |
| H3K18ac | 55 |
| H3K23ac | 55 |
| H3K27ac | 55 |
| H3K122ac | 55 |
| H4K5ac | 55 |
| H3K27me3 | 17 |
| H3K36me3 | This study |
| H3K79me1 | This study |
| H3K79me2 | This study |
| H3K4me1 | 55 |
| H3K4me3 | 17 |
| p300 | 55 |
| CBP | 55 |
| H2A.Z | This study |
| H3.3 | 56 |
| WGBS | 54 |

**Table S3:** Antibodies used in this study and published datasets

| Antibody | Company | Catalog number | Application |
| --- | --- | --- | --- |
| H3K18ac | Active Motif | 39755 | ChIP |
| H3K23ac | Millipore | 07-355 | ChIP |
| H3K27ac | Abcam | ab4729 | ChIP |
| H3K122ac | Abcam | ab33309 | ChIP |
| H4K5ac | Millipore | 07-327 | ChIP |
| H3K27me3 | Millipore | 07-449 | ChIP |
| H3K36me3 | Active Motif | 61101 | ChIP |
| H3K79me1 | Abcam | ab2886 | ChIP |
| H3K79me2 | Abcam | ab3594 | ChIP |
| H3K4me1 | Abcam | ab8895 | ChIP |
| H3K4me3 | Abcam | ab8580 | ChIP |
| p300 | Santa Cruz | sc-585X | ChIP |
| CBP | Cell Signaling Technology | 7389 | ChIP |
| H2A.Z | Abcam | ab4174 | ChIP |
| H3.3 | Millipore | 05904 | ChIP |
| H3 | Abcam | ab1791 | ChIP |
| H4 | Abcam | ab7311 | ChIP |
| IgG | Abcam | ab46540 | ChIP |
| BATF | Brookwood Biomedical | PAB4003 | Western |
| CD27 | Santa Cruz | sc-25289 | Western |
| CD70 | Santa cruz | sc-365539 | Western |

**Table S4:**
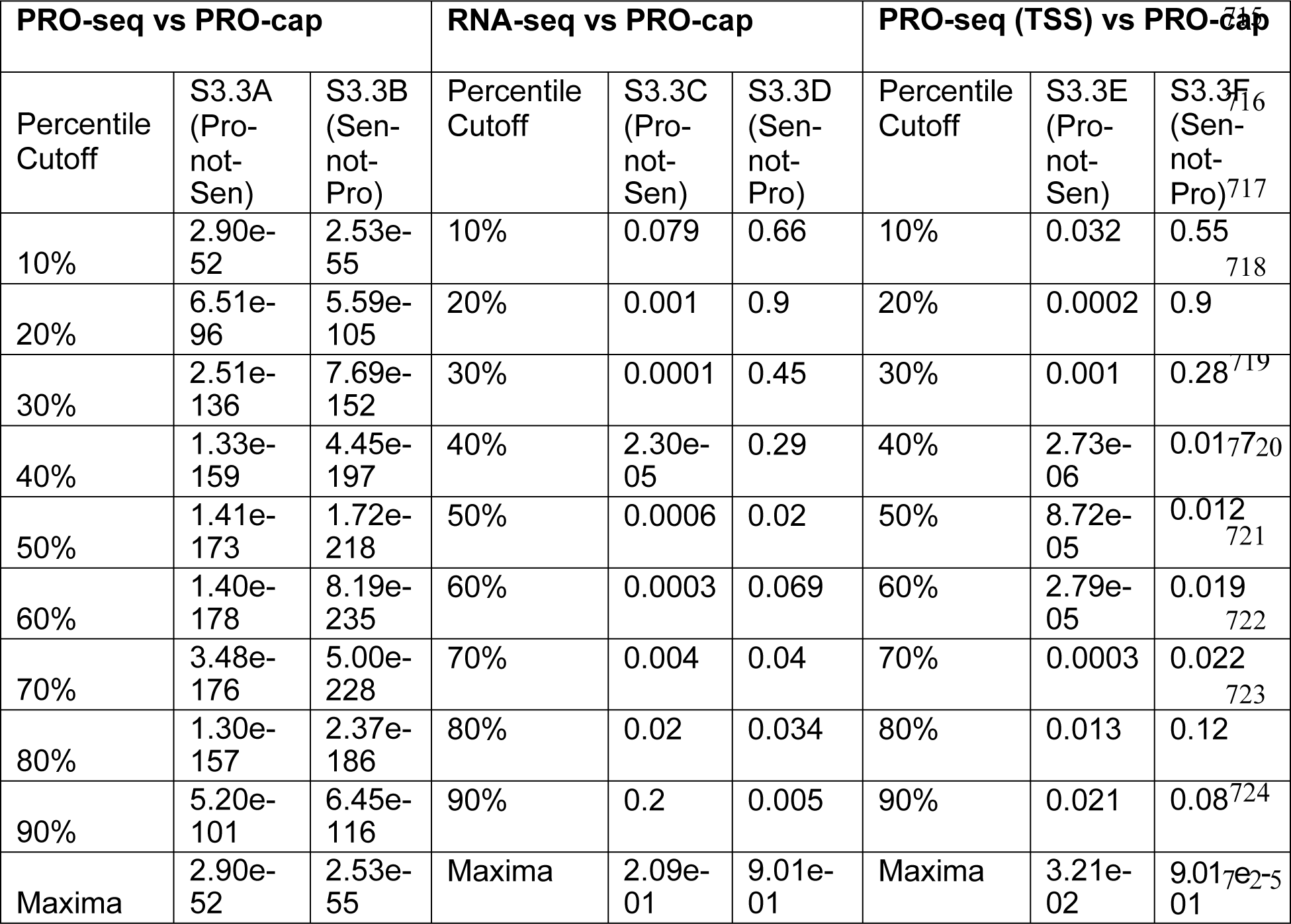
Decile-based p-values for Fig. S3.3

